# PARP1-mediated 5’ flap dynamics facilitate Okazaki fragment maturation

**DOI:** 10.64898/2026.04.27.721142

**Authors:** Guojun Shi, Yixing Wang, Yao Yan, Kejiao Li, Lingzhi Ma, Yi Lei, Yingying Wang, Nancy Manriquez, Mian Zhou, Shan Zha, Li Zheng, Binghui Shen

## Abstract

Precise regulation of enzyme recruitment during Okazaki fragment maturation (OFM) is essential for faithful and efficient lagging-strand DNA synthesis. Emerging evidence suggests that PARP1 contributes to OFM yet its specific functions remain unclear. Here, we define context-dependent functions of PARP1 during OFM. Under physiological conditions, PARP1 co-localizes with PCNA in early S phase and restrains Pol δ–PCNA– mediated strand-displacement DNA synthesis, thereby preventing the formation of long 5′ flaps, which is refractory to FEN1 cleavage. On the other hand, in LIG1-deficient cells, in which DNA nicks and unexpectedly long 5′ flaps accumulate, PARP1 promotes the recruitment of LIG3 to catalyze OF ligation and DNA2 to facilitate long 5′ flap processing. Collectively, our findings uncover previously unrecognized roles of PARP1 in regulating 5′ flap dynamics to ensure efficient OFM and cell viability.

**Highlights:**

- PARP1 plays context-dependent regulatory functions in Okazaki fragment maturation (OFM).
- PARP1 controls strand displacement DNA synthesis by the PCNA-Polδ complex to dictate generation of short over long RNA-DNA flaps during canonic OFM.
- PARP1 senses unligated Okazaki fragments in DNA Ligase 1 deficient cells and suppresses unwanted conversion of DNA nicks into 5’ flaps.
- Processing of unligated nicks or flaps by DNA ligase 3 or DNA2, respectively in LIG1 deficient cells depends on PARP1.
- PARP1 inhibitors induce synthetic lethality with DNA ligase 1 or DNA2 inhibition.

## Main

Faithful DNA replication requires efficiently and properly processing different replication intermediate structures. Of varying intermediate structures, Okazaki fragments (OFs) occurring during lagging strand DNA synthesis are the most frequent. These fragments are initiated by RNA-DNA primers synthesized by the primase-polymerase α (Pol α) complex, extended by DNA polymerase δ (Polδ), and subsequently processed to remove the RNA primers by flap endonucleases typified by flap endonuclease 1 (FEN1)(Burgers, 2009; Burgers and Kunkel, 2017). The remaining nicks between adjacent fragments are sealed primarily by DNA ligase 1 (LIG1), ensuring continuity of the lagging strand(Levin et al., 1997; Matsumoto and Kim, 1995). This process, which is named Okazaki fragment maturation (OFM), is crucial to maintain high-fidelity and efficiency of DNA synthesis (Burgers and Kunkel, 2017). Defects in Okazaki fragment processing can result in DNA nicks, replication fork collapse, genome instability, and cell death (Burgers and Kunkel, 2017).

Removal of RNA-DNA primer lies at the center of OFM. During RNA-DNA primer removal, first Pol δ catalyzes DNA strand displacement DNA synthesis (SDDS) that results in a 5’ RNA-DNA primer flap of the downstream OF(Burgers, 2009; Stith et al., 2008). If the 5’ flap is short (2-11 nt), it is effectively removed by FEN1 (Liu et al., 2004; Zheng and Shen, 2011). However, if a 5’ flap longer than 30 nt arises, ssDNA binding protein RPA may bind to the long 5’ flap and inhibit FEN1’s activity (Bae et al., 2001). In such a case, RPA recruits another 5’ flap endonuclease, DNA2, which cleaves the long 5’ flap in the middle, generating a short flap for FEN1 to cut (Bae et al., 2001).

Compared to short 5’ flaps, long 5’ flaps are more likely to form secondary structures, which may be resistant to nuclease cleavage(Nguyen et al., 2011; Zheng et al., 2020). In most cases, unprocessed 5’ flaps prevent ligation of OFs, leading to DNA single-strand breaks (SSBs) and subsequent double-strand breaks (DSBs) or collapsed replication forks, which activate the DNA damage response to induce cell cycle arrest or apoptosis. In certain genomic loci, 5’ flaps may anneal to the template strand and lead to duplication mutations or may be transformed into 3’ flaps that can form hair-pin structures or anneal and extend, generating alternative duplications with short intervening spacer sequences (Li et al., 2025; Sun et al., 2023). Due to the pathogenic consequences of long 5’ flap, the RNA-DNA primer is mostly removed by the short flap pathway. However, it is unclear about the mechanism for controlling SDDS to form short 5’ flaps that are optimal for FEN1 cleavage.

Ligation of processed OFs is the final step in OFM to produce intact lagging-strand DNA. The efficiency and accuracy of this process are crucial for genome integrity. Prolonged existence of DNA nicks may also result in DSBs or collapsed forks (Vriend and Krawczyk, 2017; Xu et al., 2025). LIG1, in complex with PCNA, is the primary enzyme for joining OFs(Blair et al., 2022; Levin et al., 2000; Tom et al., 2001). Inactivation or loss of LIG1 disrupts Okazaki fragment ligation, leading to persistent single-strand nicks at replication forks(Williams et al., 2021). It has been suggested that in mammalian cells with LIG1 deficiency, LIG3 is required for OF ligation (Arakawa et al., 2012; Arakawa and Iliakis, 2015; Kumamoto et al., 2021). On the other hand, due to the low fidelity nature of LIG3, it is normally excluded from the nick between Okazaki fragments to avoid DNA mutations. However, the mechanism for inducing LIG3 is unclear, and the consequences of activating this alternative OF ligation process are not defined.

Coordination of the recruitment and the activity of different OFM core enzymes is crucial for efficient OFM. Previous studies have shown that key steps in OFM are coordinated by PCNA (Essers et al., 2005; Matsumoto et al., 2020; Sporbert et al., 2005). PCNA interacts with Polδ, FEN1, and LIG1 to allow these core enzymes sequentially associate with OF for processing (Dovrat et al., 2014; Yan et al., 2025). PCNA first recruits Polδ to catalyze SDDS producing RNA-DNA primer 5’ flaps. Polδ is then unloaded from PCNA and FEN1 is recruited to mediate 5’ flap cleavage. Once FEN1’s job is done, it dissociates from PCNA, allows LIG1 to be recruited to join the nick. Disruption of PCNA coordination impairs OFM efficiency and fidelity (Matsumoto et al., 2020; Zheng et al., 2011; Zheng et al., 2007). In addition, protein post-translational modifications have been suggested as another important mechanism for regulating OFM processes. For instance, FEN1 arginine methylation by PRMT5, phosphorylation by CDK1, and ubiquitination by PRP19 are important for loading and unloading FEN1 to PCNA fork RNA-DNA primer removal (Guo et al., 2010; Li et al., 2018). More recently, PARP1 was found to be associated with replication forks especially when the function of FEN1 or LIG1 is deficient or inhibited in mammalian cells (Elsborg et al., 2025; Hanzlikova et al., 2018; Kumamoto et al., 2021; Yan et al., 2025). PARP1 catalyzes addition of Poly(ADP-ribose groups to protein or DNA (PARylation) to mediate protein-protein interactions or enzyme activities during DNA replication and repair processes (Hanzlikova et al., 2018; Kanev et al., 2024). The role(s) of PARP1 in regulating OFM core enzymes or other factors during OFM is unclear.

In current studies, we report that PARP1 dynamically associates with PCNA in early S phase. PARP1 is important for controlling SDDS by the Polδ-PCNA complex and PARP1 inhibition results in accumulation of 5’ flaps especially when FEN1 is inhibited. Meanwhile, LIG1 deficiency, which impairs ligation of Okazaki fragments, results in accumulation of nicks and surprisingly 5’ flaps including the long 5’ flap as well. To process these intermediate structures in LIG1 deficient cells, increased levels of PARP1 were recruited to replication forks. PARP1 in turn assist recruiting DNA2 to process long 5’ flap as well as recruit LIG3 to join the nick. PARP1 inhibition or DNA2 inhibition in LIG1 deficient cells results in significantly more DSBs and cell death than in the WT. These findings highlight important functions of PARP1 for promoting OFM and cell survival by limiting strand displacement DNA synthesis, facilitating long 5’ flap processing, and inducing alternative OF ligation when LIG1 is deficient.

## Results

### PARP1 dictates SDDS during Okazaki fragment maturation

To determine the function of PARP1 in OFM, we defined co-localization of PARP1 with PCNA, which is the scaffold protein for OFM, at different stages of S phase. It was previously shown that PCNA forms distinct focus pattern during different stages of S phase (Leonhardt et al., 2000). In early S phase, PCNA foci with relatively low intensity distributes uniformly. As S phase processes, both the number of and the intensity of PCNA foci increase. In addition, in the middle S phase, a ring of PCNA foci forms around the nuclei, and in the late S phase, PCNA foci cluster one another (Leonhardt et al., 2000). We observed that PARP1 formed foci at early S phase and they co-localized with PCNA, however, they dissociate from PCNA at late S phase (Figure 1a). In late S phase, little PARP1 foci was co-localized with PCNA foci. In early S phase, PCNA is in complex with Polδ to extend the Okazaki fragment and catalyzes SDDS to form 5’ flap for RNA-DNA primer removal (Figure 1a). We tested if PARP1 regulates SDDS. We reconstituted the SDDS assay using purified Polδ, PCNA, and PARP1 and a FAM-labeled gapped DNA substrate. We showed that the Pol δ–PCNA complex performs gap filling and strand displacement DNA synthesis, thereby generating 5′ flap intermediates. (Figure 1b). However, PARP1 on its own effectively inhibited SDDS but not gap filling, generating products with considerably shorter flaps or no flaps (Figure 1b). Inclusion of NAD⁺ to initiate PARylation reactions slightly further reduced the SDDS reaction (Figure 1b). It indicates that PARP1 physical presence is the primary mechanism in controlling Pol δ-mediated SDDS activity during OFM. It is possible that PARP1 competes with Polδ for interaction with PCNA, so as to control the processivity of Polδ or unload Polδ from the OF. Consistent with this hypothesis, we observed significantly higher levels of Polδ- PCNA complex in PARP1^-/-^ cells than in the WT cells (Figure 1c). Treatment of the cells with Olaparib, a potent PARP inhibitor (PARPi) impaired PARP1 association with PCNA and increased the PARP1-PCNA complex (Figure 1d). Nuclear extract from the cells treated with PARPi but not DMSO resulted in extensive SDDS in vitro (Figure 1e).

**Figure 1.**
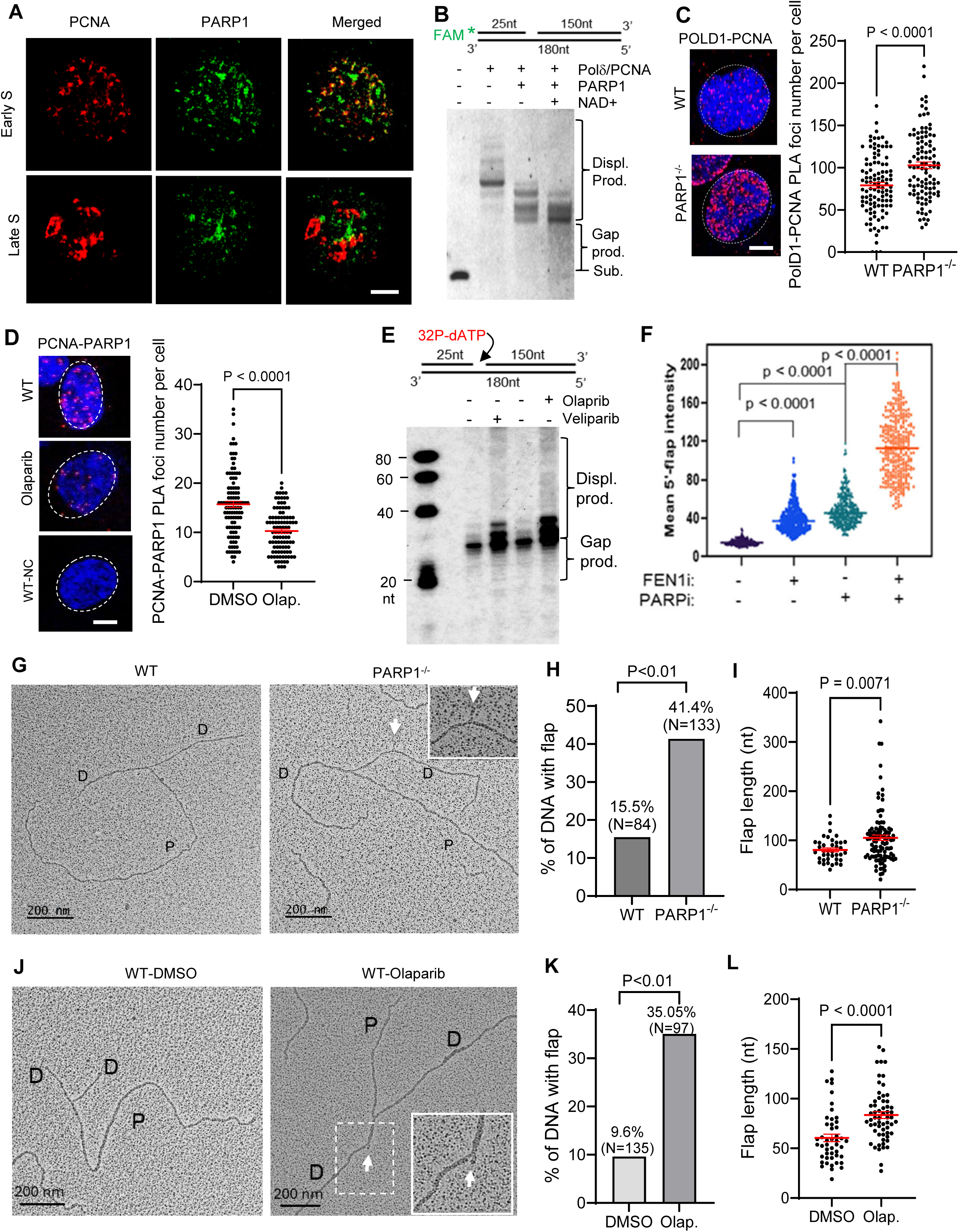
PAPR1 restrains strand displacement DNA synthesis during OFM under physiological conditions. **a.** Co-localization of PARP1 with PCNA at various early or late S phase in WT MEFs. S-phase stages were defined based on the characteristic distribution patterns of PCNA foci. **b.** Strand displacement DNA synthesis by purified Pol δ in the absence or presence of PARP1 was assayed using a FAM-labeled gapped DNA duplex. Reactions were carried out at 37 °C for 15 min. Products were analyzed by 15% denaturing PAGE. Top, schematic of gapped substrates; bottom, representative PAGE images. Substrates and reaction products are indicated. **c.** Proximity ligation assay (PLA) of POLD1 and PCNA was performed in WT and PARP1⁻/⁻ U2OS cells to assess Pol δ–PCNA complex formation. Left, representative PLA images. Red, PLA foci; blue, DAPI. Right, quantification of PLA foci per nucleus. **d.** PLA of PARP1 and PCNA was performed in MDA-MB-231 cells treated with DMSO or the PARP inhibitor olaparib (1 µM, 16 h). Left, representative PLA images. Red, PLA foci; blue, DAPI. WT MDA-MB- 231 cells incubated with a single anti-PCNA antibody were used as a negative control. Right, quantification of PLA foci in untreated and olaparib-treated nuclei. **e.** Strand displacement DNA synthesis by nuclear extracts (NEs; 1 µg per reaction) from untreated, olaparib-treated, or veliparib-treated cells was assayed using a gapped DNA duplex, [³²P]-dATP, and the remaining three deoxyribonucleotides in an ATP-free buffer to prevent DNA ligation. **f.** MDA-MB-436 cells were treated with DMSO (control), the FEN1 inhibitor LNT-1 (20 µM, 16 h), the PARP inhibitor olaparib (10 µM, 16 h), or both. 5′ flaps were labeled in situ using biotin-conjugated D181A FEN1. Flap signal intensity per nucleus was quantified using ImageJ. More than 200 nuclei per condition were analyzed (n = 2 independent experiments). P values were calculated using Student’s *t* test. **g.** Representative TEM images of replication forks in WT and PARP1^-/-^ U2OS cells. D, daughter strand; P, parental strand. Scale bars, 200 nm [500 base pairs (bp)]. White arrowheads indicate flap DNA in original or enlarged views. **h.** Percentage of replication forks containing flaps in WT and PARP1⁻/⁻ U2OS cells. Total forks analyzed: WT, n = 84; PARP1⁻/⁻, n = 133. **i.** Quantification of flap length in WT and PARP1⁻/⁻ U2OS cells. **j.** Representative TEM images of replication forks in MDA-MB-231 cells untreated or treated with olaparib. **k.** Percentage of replication forks containing flaps in untreated and olaparib-treated MDA-MB-231 cells. Total forks analyzed: untreated, n = 135; olaparib-treated, n = 97. **l.** Quantification of flap length in untreated and olaparib-treated MDA-MB-231 cells. Bars represent mean ± SEM. P values were calculated using an unpaired *t* test.

To further determine the impact of PARPi on dynamic formation and processing of 5’ flaps in human cells, we developed an in situ 5’ flap labeling protocol, in which a biotin-linked FEN1 D181A mutant-based probe (5’ flap probe) specifically binds to 5’ flaps but not nicks (Extended Data Figure S1). This technology allowed us to label and visualize 5’ flaps in single mammalian cell level under a fluorescence microscope. Considerable levels of red fluorescence signals were observed in the slide of incubation of the 5’ flap probe with MAD-MB-436 (BRCA1+) cells treated with LNT-1 (Extended Data Figure S1), which is an inhibitor of FEN1 and EXO1 (Exell et al., 2016). To prove the specificity of red fluorescence was from the 5’ flaps, the fixed LNT-1-treated cells were pre-treated with 5’ ssDNA nuclease (E. coli Rec J) or FEN1 in vitro, to cleave 5’ flaps. RecJ addition diminished or reduced the red fluorescence signal (Extended Data Figure S1, S2), demonstrating the correlation of red fluorescence signals and 5’ flap levels in the cells. In addition, we found that treatment of cells with Mimosine, which arrest cells at G1/S boundary (Krude, 1999; Kubota et al., 2014; Watson et al., 1991), significantly reduced EdU-positive cells and the flap levels in the cells (Extended Data Figure S3). It indicates that 5’ flap detected with the 5’ flap probes are predominantly associated with DNA replication. Using this approach, we determined the impact of Olaparib or/and LNT-1 on 5’ flap levels in human cells. In the absence of that PARPi or LNT-1, 5’ flaps were hardly detected (Figure 1f). LNT-1 or olaparib caused significant accumulation of 5’ flaps, and Olaparib and LNT-1 had a synergy in causing accumulation of 5’ flaps in human cells (Figure 1f).

Next, we utilized transmission electron microscopy (TEM), which is an efficient tool for detecting DNA replication intermediates, to directly visualize the presence of flap structures in human cells without or with PARP1 gene deficiency or PARP1 inhibition by its inhibitor Olaparib. We detected Y-shaped replication fork structures with one parental strand (P) and two daughter strands (D) of equal length and scored the flap structure within the daughter strand (Figure 1g). 41.4% of replication forks contained at least 1 flap structure in PARP1^-/-^ U2OS cells, compared to the WT (Figure 1h). The flap frequency in PARP1^-/-^ U2OS cells was 2.03x10^-4^ ± 2.68x10^-5^, compared to 8.48x10^-5^ ± 3.31x10^-5^ in the WT (Extended Data Figure 4a). Furthermore, the mean flap length in PARP1^-/-^ U2OS cells and the untreated control were 105.3±5.497 nt and 80.66±3.875 nt, respectively (Figure 1i). Consistently, 35.5% of replication forks contain at least 1 flap structure in Olaparib-treated MDA-MB-231 cells, compared to 9.6% in the untreated control (Figure 1j). The flap frequency in Olaparib-treated MDA-MB-231 cells was 1.61x10^-4^ ± 3.03x10^-5^ , compared to 1.55x10^-5^ ± 5.65x10^-6^ in the untreated control (Extended Data Figure 4b). The mean flap length in Olaparib-treated cells and the untreated control was 83.57±3.47 nt and 60.52 ± 3.80 nt, respectively (Figure 1k). These findings demonstrate that PARP1 is crucial in controlling the formation of 5’ flaps during OFM and PARP1 deficiency or inhibition causes accumulation of 5’ flaps including long 5’ flaps.

### PARPi exhibits synthetic lethality or growth defect with OFM genes with LIG1 as the top candidate in CRISPRi-based screening

Cancer cells frequently harbor genetic mutations such as BRCA1/2 and cause accumulation of intermediate DNA structures (Mijic et al., 2017; Schreuder et al., 2024). PARP1 plays crucial roles in sensing and/or repairing these abnormal DNA structures, therefore PARPi causes synthetical lethality (SL) or synthetic growth defect (SGD) with these genetic mutations including BRCA1/2 mutations in human cancer cells (Ashworth and Lord, 2018; Helleday, 2011). To identify new SL or SGD partners of PARPi, we carried out single cell sequencing and CRISPRi-based screening of DNA replication and repair genes (Figure 2a). The CRISPRi-based screening system covers 344 known DNA replication and repair genes (Extended Data Figure S5). If a gene deficiency causes SGD or SL, the sgRNA against this gene will be depleted during the selection process. Conversely, if gene deficiency promotes cell survival in the presence of PARPi, the sgRNA against this gene will be enriched. We observed that besides sgRNAs against genes involved in inter-crosslinking repair, sgRNAs against genes for DNA replication fork processing mainly OFM genes were significantly depleted in PARPi (Olaparib)-treated cells (Figure 2b, 2c). This suggests that OFM genes are a new category of genes that induce SL or SGD with PARPi. Of note, LIG1 was depleted to an extent similar to BRCAness genes BRCA1 or BARD1 in PARPi-treated cells (Figure 2c, 2d). As expected, sgRNAs against PARP1 were enriched, which is consistent with previous reports that loss of PARP1 gene makes cells insensitive to PARPi (Figure 2c).

**Figure 2.**
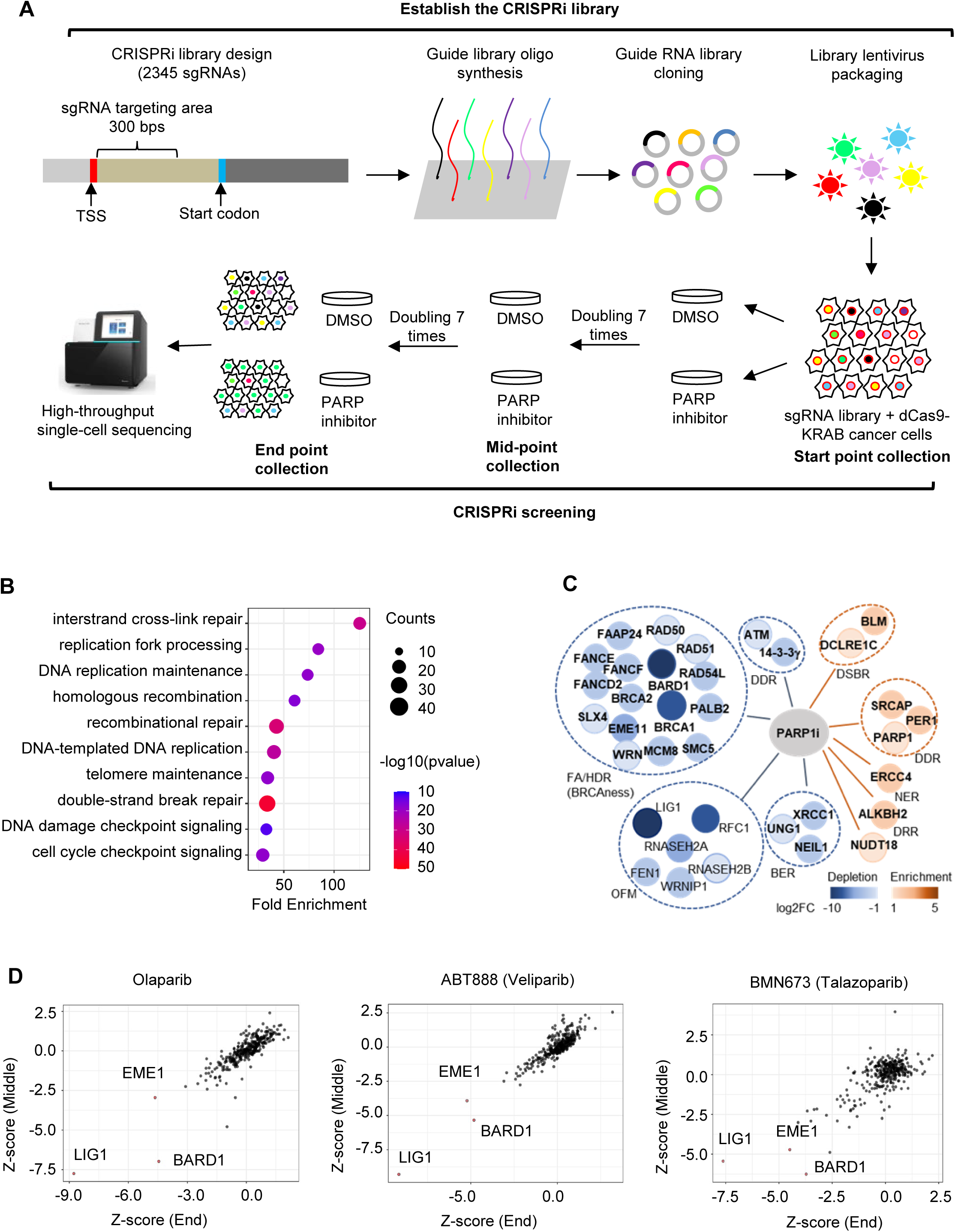
CRISPRi-based screening of genes that are depleted or enriched in PARPi-treated MDA-MB-231cells. Lentivirus-based CRISPRi library against 344 known DNA replication and repair genes (7 different sgRNA for each gene) were infected with MDB-MA-231 cells that are cultured in the absence or presence of Olaparib (10 µM), Veliparib (50 μM), or BMN673 (5 nM) (n = 3 for each group). Cells were collected at 0, 7, and 14 doublings and subjected to single-cell based DNA sequencing to determine the copy number of a specific sgRNA in different cell populations. **a.** Diagram outline CRISPRi library construction, oligo synthesis, drug selection, and single cell sequencing in the CRISPRi-based screening. **b.** Pathway enrichment analysis of genes with negative Z-scores (Z < 0) across PARP inhibitor treatments. **c.** Depleted or enriched genes in different DNA replication and repair pathways in olaparib-treated cells. **d.** Correlation of CRISPRi Z-scores between mid- point and end-point time points for individual PARP inhibitors.

These results suggest a crucial function of PARP1 in mediating alternative OF ligation processes, which underly the SL phenotype between LIG1 deficiency and PARPi.

### LIG1 deficiency and PARP1 inhibition synergistically cause accumulation of flaps, SSBs, and DSBs

Deficiency in LIG1 caused accumulation of un-ligated Okazaki fragments (Smith and Whitehouse, 2012; Soza et al., 2009). Theoretically, the unligated nicks may be processed into ssDNA gaps by exonucleases, flaps by polymerases or helicases, or DSBs by endonucleases. We constructed LIG1 knockdown cells using the CRISPRi strategy, which effectively reduced the LIG1 protein level (Extended Data Figure S6) and used them to analyze the molecular alterations in the absence or presence of PARP inhibitors. We used TEM to directly assess replication intermediates in WT or LIG1 knockdown MDA-MB-231 cells. Y-shaped replication fork structures with two daughter strands of equal length were detected in both WT and LIG1 knockdown cells (Figure 3a). Notably, short DNA flaps were detected in daughter strands of 47.1% replication forks in LIG1 knockdown cells, compared to only 9.6% in WT cells (Figure 3b). Olaparib treatment further increased the percentage of flap-containing replication forks to 52.2% in LIG1 knockdown cells (Figure 3b). Flaps were also detected in linear DNA regions in LIG1 knockdown cells but not in the WT cells without Olaparib treatment (Extended Data Figure S7). On the other hand, flaps were detected in the linear DNA regions in Olaparib-treated WT and LIG1 knockdown cells (Extended Data Figure S7). The flap frequency in LIG1 knockdown cells was 2.37x10^-4^ ± 6.30x10^-5^, compared to 1.55x10^-5^ ± 5.65x10^-6^ in the WT (Figure 3c). The flap frequency in Olaparib-treated WT and LIG1 knockdown cells was significantly increased to 1.61x10^-4^ ± 3.03x10^-5^ and 5.36x10^-4^ ± 1.22x10^-4^, respectively (Figure 3c). Furthermore, the mean flap length in LIG1 knockdown was 85.14±3.898 nt, compared to 60.52±3.799 nt in the WT, and Olaparib treatment significantly increased the flap length in both WT and LIG1 knockdown cells (Figure 3d). Consistently, BrdU alkaline comet assay revealed that LIG1 knockdown cells accumulated significantly more SSBs at the nascent DNA strand than the WT cells (Figure 3e-3f). Neutral and alkaline comet assays revealed that LIG1 knockdown cells had a significant increase in DSB levels, compared to the WT (Figure 3g-3h and Extended Data Figure S8). Furthermore, LIG1 knockdown and Olaparib treatment had a synergy in inducing SSBs and DSBs (Figure 3e-3h). These findings demonstrate that PARP1 plays important roles in processing flaps, repairing nicks, and avoiding DSB accumulation in LIG1 deficient cells.

**Figure 3.**
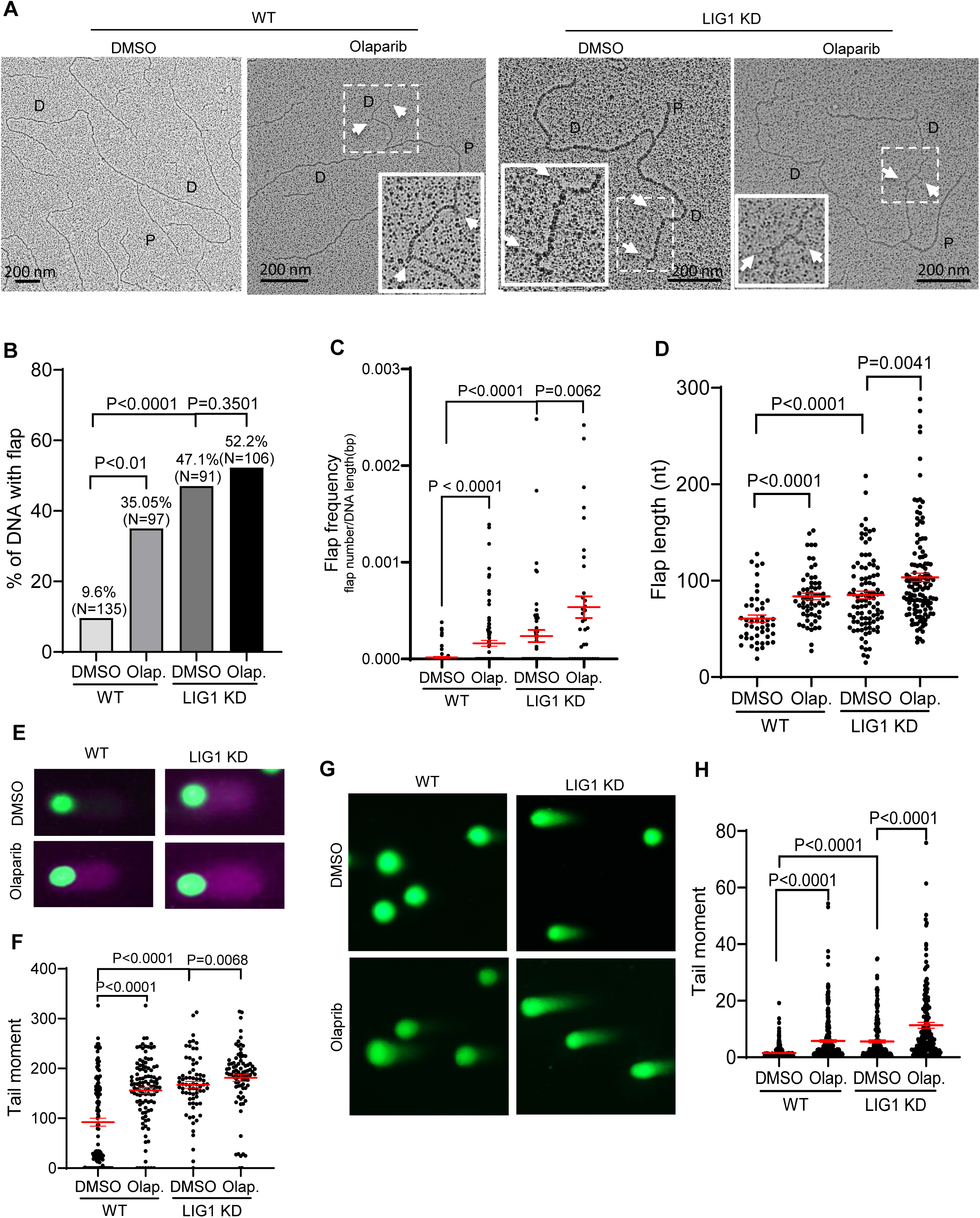
LIG1 deficiency and Olaparib have synergy in accumulation of OFM related flaps, SSBs, and DSBs. **a.** Representative TEM images of replication forks in WT and LIG1 knockdown MDA-MB-231 cells untreated or treated with Olaparib (1 μM, 16 h). D, daughter strand; P, parental strand. Scale bars, 200 nm [500 base pairs (bp)]. White arrowheads indicate flap DNA in the original or enlarged view. **b-d.** Quantification of the percentage of replication forks containing flaps (B), flap frequency (C), and flap length (D) in WT and LIG1 knockdown MDA-MB-231 cells untreated or treated with olaparib (1 µM, 16 h). Flap frequency was calculated as the number of flaps divided by the length of the analyzed DNA fragment. **e. and f**. Alkaline BrdU comet assays in WT and LIG1 knockdown MDA-MB-231 cells untreated or treated with olaparib (1 µM, 16 h). Cells were labeled with BrdU (20 µM, 20 min) prior to alkaline comet analysis. **g. and h.** Neutral comet assays in WT and LIG1 knockdown MDA-MB-231 cells untreated or treated with Olaparib (1 μM, 16 h). Panels E. and G. show representative comet images; panels F and H are tail moments of individual nucleus. For each sample, at least 50 comets were quantified. Median tail moments are indicated on the graph. P values were calculated by the two-sided student’s t-test.

### PARP1 Inhibition distinctly impacts on DNA Damage response, genome stability, and cell survival in LIG1 knockdown Cells

Next, we determined the impact of PARP1 inhibition on DNA damage response, cell proliferation, and survival, and genome integrity of WT and LIG1 knockdown cells. We observed that LIG1 knockdown cells exhibited a significant increase in the level of γH2AX foci, a well-established marker of DNA damage (Kuo and Yang, 2008; Mah et al., 2010), compared to WT cells (Figure 4a and 4b). To evaluate of the impact of PARP1 inhibition in DNA damage response in the WT cells and LIG1 knockdown cells, we treated the cells with olaparib and assessed the levels of γH2AX. Olaparib led to elevated levels of γH2AX foci in both WT and LIG1 knockdown cells in a dose-dependent manner, but it affected LIG1 knockdown cells more that the WT cells (Figure 4a, 4b). Olaparib at 0.1 μM significantly increased the number of γH2AX foci in LIG1 knockdown cells but not in the WT cells (Figure 4a, 4b). Consistently, western blot analysis showed that LIG1 knockdown cells had higher level of γH2AX than the WT cells in the absence or presence of varying concentrations of olaparib (Figure 4c).

**Figure 4.**
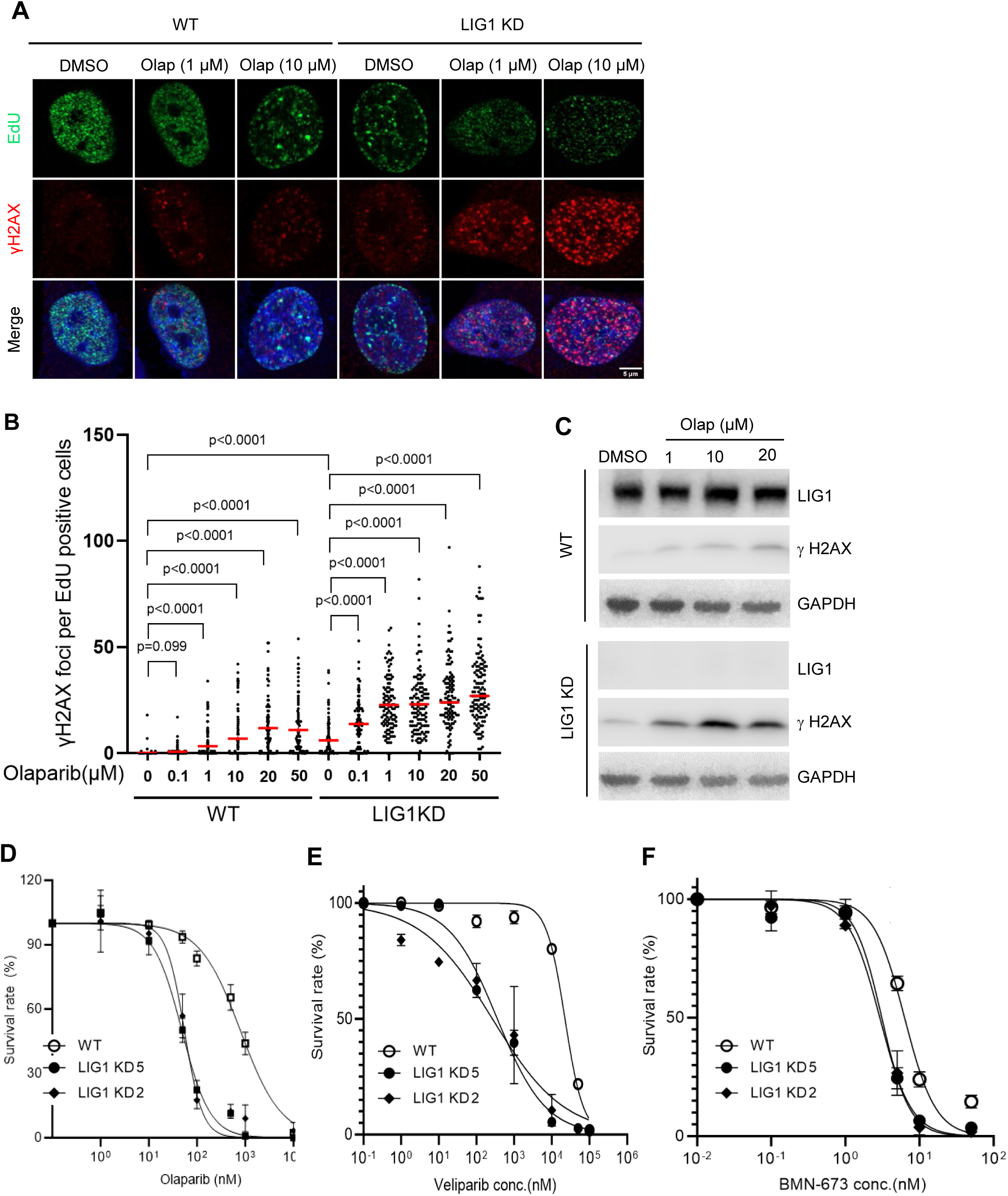
LIG1 deficiency sensitizes cells to olaparib. **a**. and **b.** γH2AX foci in EdU- positive WT and LIG1 knockdown MDA-MB-231 cells untreated or treated with Olaparib (1 μM, 16 h). Cells were labeled with EdU (10 μM, 30 m). Incorporated EdU was detected with the CLICK assay and γH2AX IF was carried out using a specific antibody against phosphorylated H2AX (S139) (γH2AX). Panel A shows representative staining images of EdU (green), γH2AX (red), and nucleus (blue). Panel B is quantification of γH2AX foci in EdU positive cells in different samples. Median levels are indicated by red lines on the graph. P values were calculated by the two-sided student’s t-test. **c.** Immunoblot analysis of total γH2AX in WT and LIG1 knockdown MDA-MB-231 cells treated with increasing concentrations of olaparib (0, 1, 10, or 20 µM) for 16 h. GAPDH served as a loading control. **d-f.** Sensitivity of WT and LIG1 knockdown MDA-MB-231 cells to olaparib (**d**), veliparib (**e**), or talazoparib (BMN-673; **f**). Parental (WT) cells and two independent LIG1 knockdown cell lines generated using distinct sgRNAs were analyzed. Data represent mean ± SD from four independent experiments.

Meanwhile, we observed that LIG1 knockdown cells had significantly slower cell growth rate than the WT (Extended Data Figure S9). Cell proliferation assays indicated that the doubling time for LIG1 knockdown cells was 29.7 ± 0.6 h, compared to 22.7 ± 0.1 h for the cells. Furthermore, PARP inhibitors hindered the cell growth of LIG1 knockdown cells but not the WT cells. In the presence of 1 μM Olaparib, 10 μM veliparib, 10 nM Talazoparib, the double time of LIG1 knockdown cells were 104.7 ± 22.5 h, 107.1 ± 12.5 h, 71.7 ± 4.6 h, respectively, compared to 25.2 ± 0.2 h, 24.0 ± 0.2 h, 27.7 ± 0.3 h, respectively for the WT cells significantly increased. Consistently, cell viability assay showed that LIG1 knockdown cells were significantly sensitive to different PARP inhibitors than the WT cells (Figure 4d). These findings support the CRISPRi-based screening result showing that LIG1 deficiency and PARP inhibition results in SL.

Next, we sought to define the frequency and spectra of mutations in WT and LIG1 knockdown cells untreated or treated with PARPi (Olaparib). Genomic DNA was isolated from different cells and whole-exome sequencing (WES) was performed. 90.4 somatic SNVs, 22.6 somatic small insertions or deletions (Indels), and 1.7 duplications per million base pairs were detected in LIG1 knockdown cells, compared to 22.8 SNVs, 2.4 Indels, and 1.5 duplications in the WT cells (Figure 5a-5c). LIG1 deficiency did not drastically change the mutation signature of cells. In both the WT and LIG1 knockdown cells, C>T and T>C were the most frequent SNVs (Figure 5d). Mutation cluster analysis revealed that LIG1 knockdown-induced mutations displayed the DBS mutation cluster (Figure 5e), which is linked to APOBEC3-mediated mutagenesis (Alexandrov et al., 2013), and Kataegis mutation cluster (Figure 5e), which is linked to HDR-mediated DNA double strand break repair (Roberts et al., 2012; Taylor et al., 2013).

**Figure 5.**
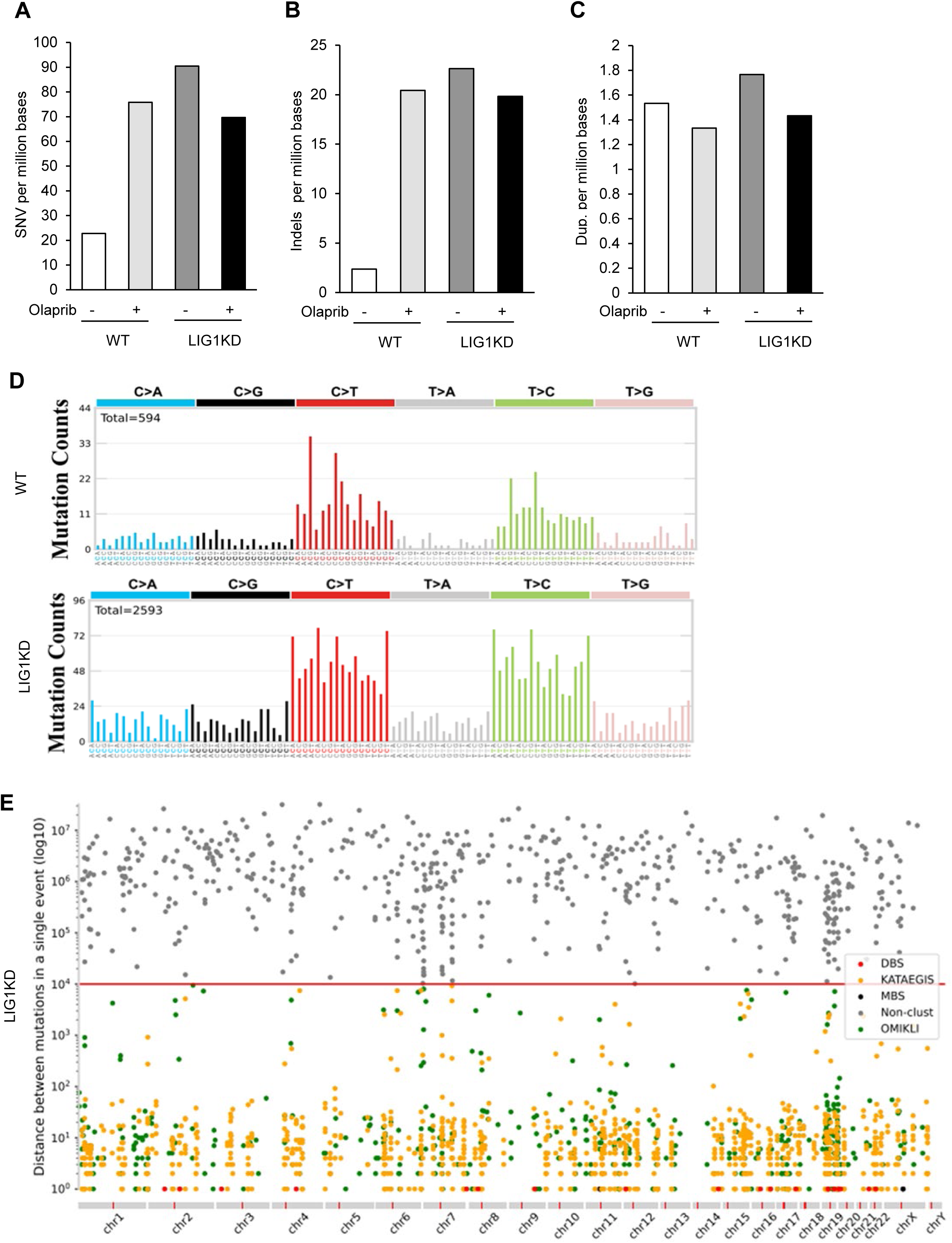
LIG1 deficiency and Olaparib induces DNA mutations. **a-c**. Whole-exome sequencing (WES) was performed to determine the frequency of single-nucleotide variants (SNVs) (A), small insertions and deletions (indels; **b**), and duplications (**c**) in WT and LIG1 knockdown MDA-MB-231 cells untreated or treated with olaparib. **d.** Mutational signature analysis in WT and LIG1 knockdown MDA-MB-231 cells. Base substitutions (C>A, C>G, C>T, T>A, T>C, and T>G) were categorized according to the 5′ and 3′ flanking nucleotide context of the mutated base. **e.** Mutation cluster analysis in WT or LIG1 knockdown MDA-MB-231 cells. Inter-mutation distances were calculated to define clustered mutations, including kataegis (break-induced clusters; orange), omikli (APOBEC3-associated clusters with short intermutation distances; green), doublet base substitutions (DBS; red), and multibase substitutions (MBS; black). All remaining mutations were classified as nonclustered.

### LIG1 deficiency induces PARP1-dependent recruitment of LIG3 and DNA2 for OFM

The fact that LIG1 knockout or knockdown cells are viable and proliferating suggests that LIG1 deficient cells activate alternative ways to process long flaps, nicks, and gaps. We hypothesize that PARP1 senses these OFM intermediate structures and induce alternative OFM processes in LIG1 knockdown cells to support cell survival. Supporting this hypothesis, we observed that PARP1 recruitment to replication forks was significantly increased in LIG1 knockdown cells or cells treated with the LIG1 inhibitor (Howes et al., 2017) as shown in the EdU-PLA assays (Figure 6a, 6b and Extended Data Figure S10a, S10b). In addition, PCNA-PARP1 PLA assay showed that EdU-positive LIG1 knockdown cells had more PARP1-PCNA complex than EdU-positive WT cells (Figure 6c, 6d).

**Figure 6.**
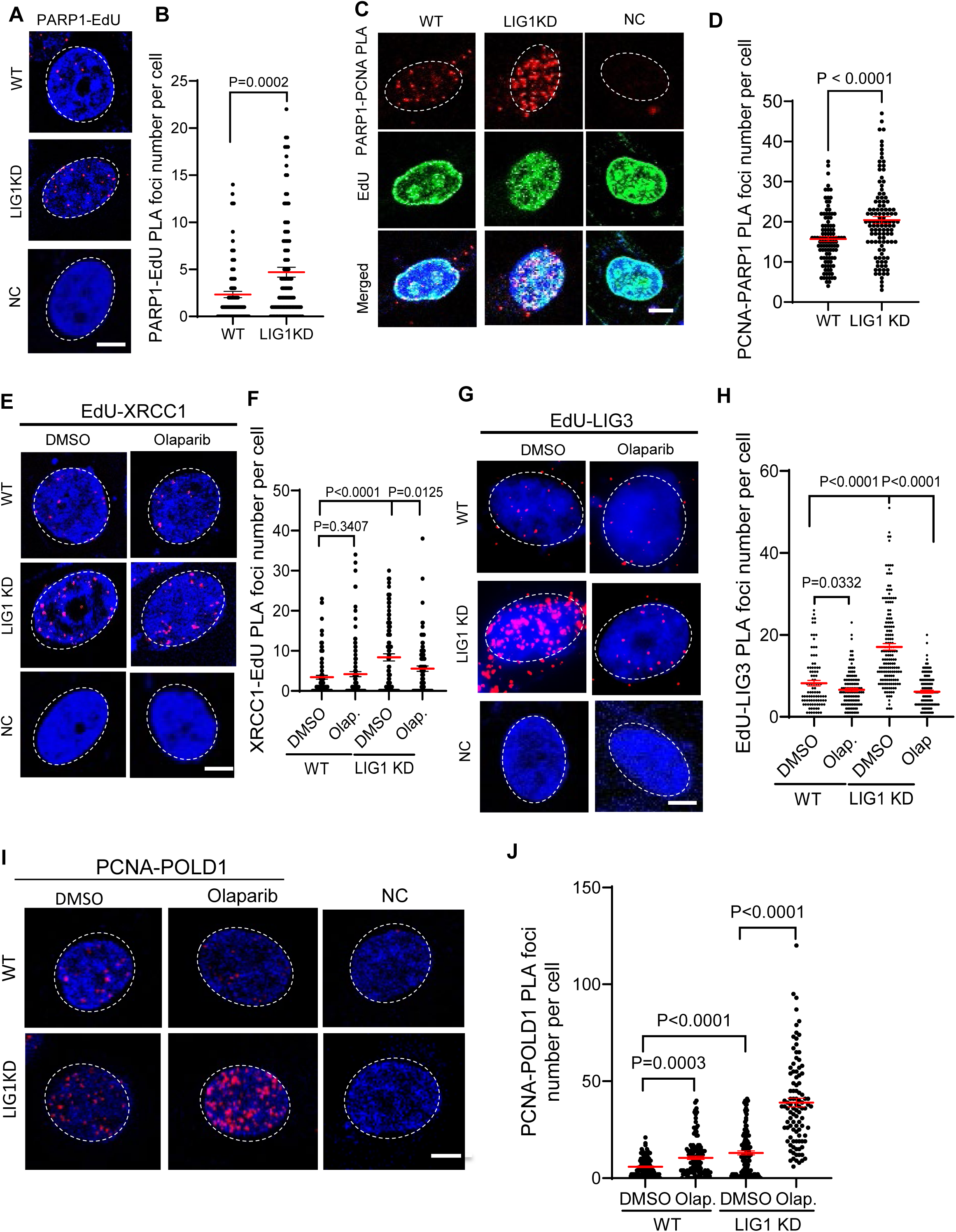
LIG1 deficiency results in additional PARP1 binding to replication forks to induce PARP1-dependent alternative OFM processes. **a**. and **b.** PARP1-EdU PLA to analyze nascent DNA-associated PARP1 in WT and LIG1 knockdown MDA-MB-231 cells. **(a)** Representative PLA images. Red, PLA foci; blue, DAPI. WT MDA-MB-231 cells incubated with a single anti-PCNA antibody were used as a negative control. Scale bar, 5 µm. **(b)** Quantification of PARP1–EdU PLA foci per nucleus. **c. and d.** PLA analysis of the PCNA–PARP1 complex in replicating WT and LIG1 knockdown (KD) MDA-MB-231 cells. Cells were labeled with EdU (10 µM, 30 min) to identify replicating cells, and PCNA-associated PARP1 was analyzed by PCNA–PARP1 PLA in EdU- positive nuclei. Nuclei were counterstained with DAPI (blue). WT MDA-MB-231 cells incubated with a single anti-PCNA antibody served as the negative control. **(c)** Representative images. **(d)** Quantification of PCNA–PARP1 PLA foci. **e-h.** PLA to analyze nascent DNA-associated XRCC1 (E, F) or LIG3 (G, H) or to analyze the PCNA-Polδ complex (I, J) in WT or LIG1 knockdown MDA-MB-231 cells untreated or treated with Olaparib (1 μM, 16h). In EdU-XRCC1 or EdU-LIG3 PLA, cells were labeled with EdU (10 μM, 30 m) prior PLA. Panels E, G, I show representative EdU-XRCC1, EdU-LIG3, or PCNA-POLD1 PLA images, respectively and panels F, H, J show corresponding PLA foci number per nucleus in different samples. Median PLA foci levels are indicated with red lines on the graph. P values were calculated by the two- sided student’s t-test.

We propose that increased presence of PARP1 at the replication forks could induce recruitment of enzymes to process OFM intermediates accumulated due to LIG1 deficiency. DNA ligase 3 (LIG3) has been proposed to compensate for LIG1 function. Consistently, we observed that LIG1 knockdown cells had significantly more XRCC1 and LIG3 that were associated with nascent DNA than WT cells (Figure 6e-6h). We also detected significantly more PCNA-colocalized LIG3 in LIG1 knockdown cells than in WT cells (Extended Data S11a, S11b). To determine whether LIG3 recruitment in LIG1 knockdown cells was dependent on PARP1, we treated the cells with Olaparib. We found that Olaparib treatments inhibited LIG1 knockdown-induced XRCC1 or LIG3 recruitment to replication forks (Figure 6e-6h). Consistently, Olaparib treatments suppress LIG3 co-localization with PCNA in LIG1 knockdown cells (Extended Data Figure S11a, S11b). At the same time, the level of PCNA-colocalized Polδ in LIG1 knockdown cells was significantly higher in the LIG1 knockdown or inhibited cells than in the WT (Figure 6i, 6j, Extended Data S12a, S12b). Olaparib treatment of LIG1 knockdown cells resulted in a further drastic increase in the PCNA-colocalized Polδ (Figure 6i, 6j).

The existence of long 5’ flaps in LIG1 knockdown cells suggests that DNA2-mediated long flap processing is needed for OFM in the mutant cells. To test this hypothesis, we analyzed DNA2 recruitment to replication forks and its interaction with PCNA in LIG1 knockdown cells. We observed that significantly more DNA2 was recruited to nascent DNA in LIG1 knockdown cells than in the WT cells (Figure 7a, 7b). The level of PCNA-colocalized DNA2 in LIG1 knockdown cells was also significantly higher in the LIG1 knockdown than in the WT (Figure 7c, 7d). While Olaparib treatment of LIG1 knockdown cells only slightly reduced the EdU-associated DNA2 (Figure 7a, 7b), it greatly inhibited co-localization of DNA2 with PCNA (Figure 7c, 7d). It suggests that PARP1 assists DNA2 interaction with PCNA to process long 5’ flaps arisen in LIG1 knockdown cells, although the recruitment of DNA2 is induced by other regulatory protein such as ssDNA binding protein RPA.

**Figure 7.**
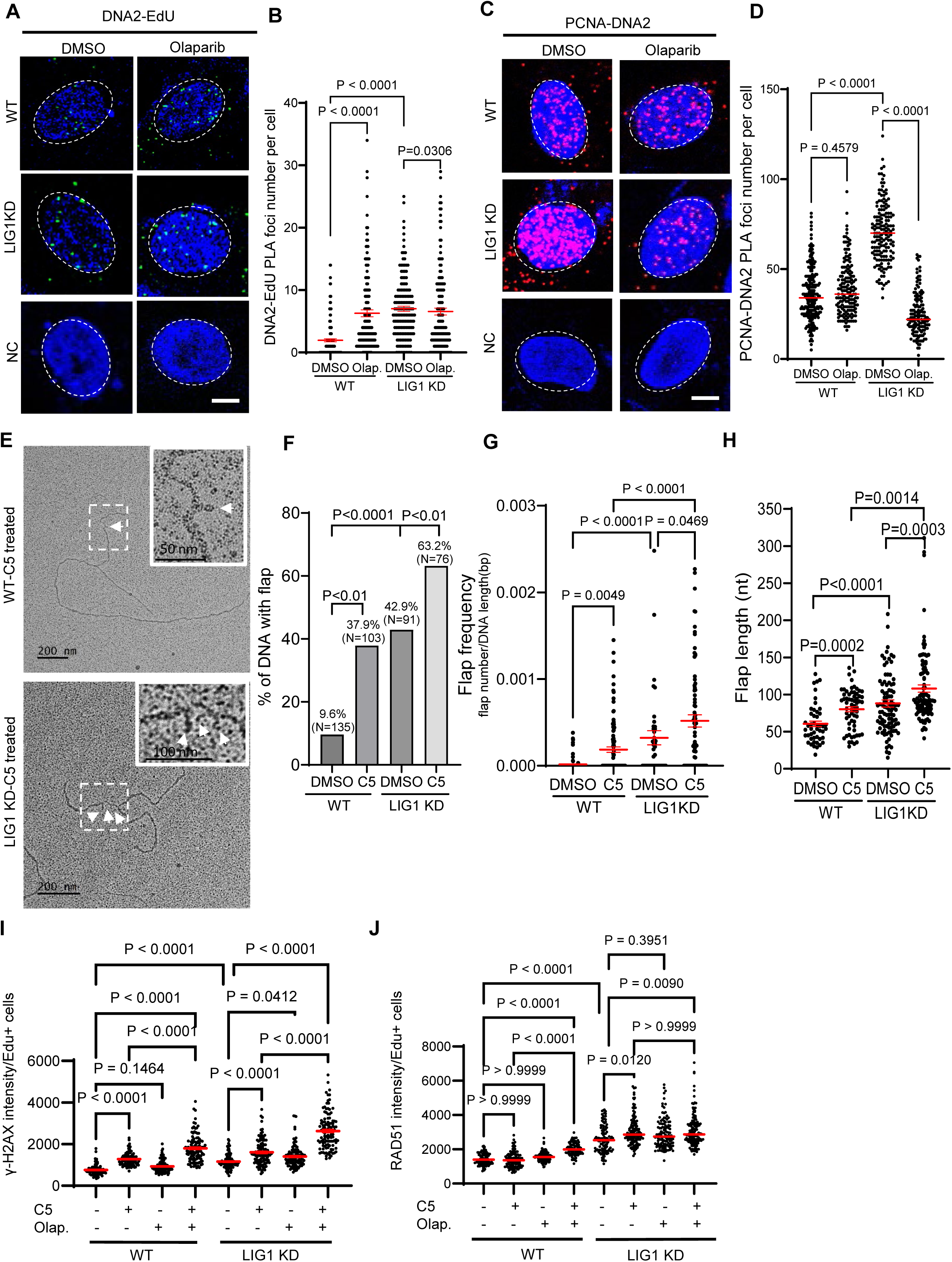
LIG1 deficiency induces a PARP1-dependent DNA2-mediated long flap processing pathway during Okazaki fragment maturation and promotes cell survival. **a-d**. PLA to analyze nascent DNA-associated DNA2 (A, B) or the PCNA- DNA2 complex (C, D) in WT and LIG1 knockdown MDA-MB-231 cells untreated or treated with Olaparib (1 μM, 16h). In EdU-DNA2 PLA, cells were labeled with EdU (10 μM, 30 m) prior to PLA. (A, C) Representative EdU–DNA2 and PCNA–DNA2 PLA images, respectively. (B, D) Quantification of PLA foci per nucleus under the indicated conditions. Median values are indicated by red lines. P values were calculated using a two-sided Student’s t test. **e.** Representative TEM images of replication forks in WT and LIG1 knockdown MDA-MB-231 cells untreated or treated with DNA2 inhibitor C5 (100 μM, 16 h). D, daughter strand; P, parental strand. Scale bars, 200 nm [500 base pairs (bp)]. White arrowheads indicate flap DNA in the original or enlarged view. **f-h.** Quantification of the percentage of replication forks containing flaps (F), flap frequency (G), and flap length (H) in WT and LIG1 knockdown MDA-MB-231 cells untreated or treated with C5. Flap frequency was calculated as the number of flaps divided by the length of the analyzed fork DNA fragment. **i. and j.** γH2AX (panel I) or RAD51 (panel J) foci in EdU-positive WT and LIG1 knockdown (LIG1 KD) MDA-MB-231 cells untreated or treated with C5 (100 μM, 16 h). Cells were labeled with EdU (10 µM, 30 min) prior to analysis. EdU incorporation was detected by Click chemistry, and γH2AX or RAD51 was detected by immunofluorescence. The number of γH2AX foci in EdU-positive cells was quantified using Image J. Median values are indicated by red lines. P values were calculated using a two-sided Student’s t test.

To test if DNA2 is crucial for processing long 5’ flaps in LIG1 knockdown cells, we inhibited DNA2 by C5 and assessed the level and length of flap structure. In the presence of C5, the flap frequency and length in LIG1 knockdown cells were significantly higher and longer than in WT cells (Figure 7e-7h). Consistently, C5 treatment significantly enhanced the level of γH2AX or RAD51 foci in LIG1 knockdown cells than in WT cells (Figure 7i, 7j, Extended Data Figure S13), suggesting that DNA2 inhibition and LIG1 deficiency have a synergy in inducing DNA damage. Furthermore, cell survival assays showed that LIG1 knockdown cells were significantly more sensitive to C5 than WT cells, especially in the presence of Olaparib (Extended Data Figure S14).

## Discussion

Our current studies define molecular mechanisms of PARP1 in regulating OFM in mammalian cells both of normal physiology or under stress due to LIG1 deficiency. We demonstrate that an important function of PARP1 during canonic DNA replication is to dictate the pathway for removal of RNA primer removal, which lies at the center of OFM and is crucial for cell survival. In mammalian cells, the RNA-DNA primer is removed via sequential action of relies on highly coordinated enzymatic reactions of Polδ catalyzed SDDS producing 5’ flaps and subsequent nuclease-mediated 5’ flap cleavage. The RNA-DNA 5’ flap shorter than 35 nucleotides can be effectively removed by FEN1 alone. However, 5’ flap longer than 35 nucleotides will be tightly bound by RPA, which inhibits FEN1’s nuclease activity. Meanwhile, RPA recruits DNA2 nuclease, which cut the long 5’ flap at the middle to convert the long 5’ flap into a short one for FEN1 to cleave (Bae et al., 2001). Long 5’ flaps have higher potential to form secondary structures leading to duplication or alternative duplication mutations (Sun et al., 2023). They may also induce DNA damage response (Nguyen et al., 2011; Zheng et al., 2020). It is generally accepted that FEN1-mediated short flap cleavage is the primary pathway to remove RNA-DNA primer (Zheng et al., 2020; Zheng and Shen, 2011). However, it is unknown about the regulator that controls Polδ catalyzed SDDS so as to produce short 5’ flaps, which are ideal substrates for FEN1. Our biochemical and cellular data provide convincing evidence supporting that PARP1 is crucial for limiting the length of 5’ flaps during OFM. PARP1 was previously shown to be localized to replication forks when OFM is disturbed. We discovered that PARP1 dynamically interacts with the OFM platform protein PCNA under normal physiology. It co-localized with PCNA in early S phase, when OF extension and SDDS occur. While the presence of PARP1 does not affect OF extension, it controls the SDDS activity of Polδ, allowing short flap formation. One possible mechanism is that PARP1 promotes Polδ dissociated from PCNA when short 5’ flap forms, because PARP1 knockout cells have significantly more PCNA-Polδ complex than WT cells. Intriguingly, we found that PARP1 inhibitor Olaparib affects the PARP1-PCNA interaction and enhances SDDS, promoting 5’ flap formation in vitro and in vivo.

We discover that PARP1 is unloaded from PCNA in middle or late S phase, when processed OFs are ligated into an intact lagging strand DNA. This step is important to avoid unwanted error prone OF ligation. While yeast relies on a single DNA ligase (Cdc9) for all ligation steps (Willer et al., 1999), mammalian cells have LIG1, LIG3 and LIG4 (Sallmyr et al., 2020). During canonic OFM in mammalian cells, OF ligation is predominantly performed by DNA ligase 1 (LIG1), which catalyzes DNA ligation of high fidelity(Williams et al., 2021). DNA nicks with nearby DNA mutations or small loops are not ligated by LIG1. On the other hand, LIG3, which has long been thought to help DNA ligation including OF when LIG1 is absent, is a DNA ligase of low fidelity. We recently showed that LIG3 can ligate nick DNA substrates with a DNA mismatch 2 nt away from the nick site. Furthermore, LIG3 can ligate 5’ flap with the upstream primer via the micro-homology DNA sequences (Yan et al., 2025). Thus, avoidance of LIG3 at replication forks to compete with LIG1 is crucial for maintaining faithful OF ligation. Because PARP1 could potentially induce recruitment of LIG3 to replication forks, its dissociation from PCNA at late S phase is important for preventing unwanted LIG3 mediated error prone OF ligation.

Genetic deficiency or chemical inhibition of LIG1 impairs canonic OF ligation, leading to induction of PARP1-dependent alternative OFM processes for survival (Kumamoto et al., 2021). LIG1 deficiency leads to retaining PARP1 at replication forks or recruiting more PARP1 to replication forks. Such molecular change is crucial for completing OFM in LIG1 deficient/inhibited cells and supporting cell survival via different mechanisms.

The presence of PARP1 at unligated nick between OFs induces recruitment of LIG3, which catalyzes OF ligation. In case that OF ligation fails, the nick DNA may be improperly processed. One scenario is re-association of DNA polymerases such as Polδ, which catalyzes SDDS to convert the nick into 5’ flaps. We show that PARP1 is important in preventing such molecular events, as PARP1 inhibition increases Polδ association with nascent DNA nicks and PCNA and results in significantly higher flaps in LIG1 knockdown cells than in WT cells. Nevertheless, considerable number of unligated nicks are converted into 5’ flaps especially long 5’ flaps in LIG1 deficient cells. The presence of the long 5’ flap induces recruitment of DNA2 to replication forks. This process depends on also PARP1, which directly binds both PCNA and DNA2. It is possible that PARP1 assists DNA2 interaction with PCNA, as PARP1 inhibition suppresses formation of the PCNA-DNA2 complex in LIG1 knockdown cells.

These findings are consistent with previous studies indicating that PARP1 acts as a key DNA damage sensor and mediator of chromatin remodeling through poly(ADP-ribosylation (PARylation) (Elsborg et al., 2025; Saldanha et al., 2024). PARP1 catalyzes the PARylation of chromatin-associated proteins or creates a docking platform for DNA repair factors (Ortega et al., 2025). During OFM in LIG1 deficient cells, PARP1 recruits LIG3 and DNA2 for inducing alternative OFM processes. Previous structural studies have demonstrated that LIG3 interacts with XRCC1 via its BRCT domain, and that XRCC1 itself is a well-known PAR-binding protein. PARylation by PARP1, therefore, likely facilitates XRCC1-LIG3 complex recruitment to sites of OF nicks (Kanev et al., 2024). Future studies are needed to clarify whether PARP1-dependent DNA2 recruitment requires XRCC1 PARylation.

When PARP1 is inhibited in LIG1 deficient cells, these alternative OFM pathways collapse, it results in excessive DNA damage including the lethal DSBs as indicated by the Comet tails and significantly elevated γH2AX levels. Consequently, LIG1 knockdown cells were especially sensitive to different PARP inhibitors, especially when DNA2 is also inhibited. The SL between LIG1 and PARP inhibitors represents. The accumulation of replicative SSBs including nicks and flaps due to LIG1 deficiency and PARP inhibitors represents a new mechanism to induce SL to treat human cancers. From a broader perspective, our findings illustrate the redundancy and plasticity of the OFM machinery in mammalian cells, where LIG3 and DNA2 assisted by PARP1, serves as emergency ligase and nuclease to support replication fork progression when OFM is impaired as in LIG1 deficient cells. This is in contrast to the budding yeast Saccharomyces cerevisiae, which relies on a single ligase (Cdc9) and lacks such compensatory flexibility(Willer et al., 1999).The evolutionary expansion of the ligase family in metazoans may reflect the need to maintain genome stability under diverse physiological and genotoxic conditions. Importantly, these findings have translational relevance. Tumors with LIG1 mutations, downregulation, or epigenetic silencing may exhibit heightened sensitivity to PARP inhibitors due to the collapse of both primary and backup ligation pathways. Thus, LIG1 status may represent a novel biomarker for predicting PARP inhibitor response beyond the current BRCA-deficiency paradigm.

## Material and Methods

### CRISPR library and single sgRNA cloning

Guide RNA design and cloning were performed as previously described (Mattson et al., 2024). sgRNA oligonucleotides were generated either by high-throughput microarray synthesis (CustomArray; for pooled library construction) or by individual synthesis (IDT; for single sgRNA experiments). Oligos were cloned into the ipUSEPR lentiviral sgRNA expression vector, which features an hU6 promoter–driven sgRNA cassette co-expressing EF-1α–driven red fluorescent protein (RFP) and a puromycin-resistance marker, using BsmBI restriction sites (NEB). The sgRNA sequences were designed using the BROAD Institute Genetic Perturbation Platform – CRISPick. For the CRISPRi library for genome stability genes (current study), 2,345 sgRNAs targeting 328 genes encoding DNA replication and repair proteins were selected (seven sgRNAs per gene). Each sgRNA was designed to target the TSS region for each gene to inhibit the transcription. All cloned libraries underwent quality control by NextSeq sequencing, ensuring that ≥90% of sgRNAs achieved at least 10 reads per million.

### Data processing and gene essentiality scores

The sgRNA read count files were computed from the raw CRISPR fastq files, for each experiment, using the ‘count’ function from the MAGeCK software (Li et al., 2014). The gene scores (normZ values) were estimated from the read count files using the DrugZ algorithm (Colic et al., 2019), testing the drug treated replicates against their respective DMSO-treated replicates. Data quality was assessed using standard diagnostics generated by MAGeCK, including sgRNA representation, sequencing depth, and read count distributions, to ensure adequate library coverage and consistency across replicates. Only datasets passing these quality control criteria were used for DrugZ analysis and downstream analyses.

### Pathway enrichment analysis

Gene Ontology (GO) Biological Process (BP) enrichment analysis was performed using genes with negative DrugZ normalized Z-scores (NormZ < 0) across all PARP inhibitor treatments. Enrichment analysis was conducted against the background of all genes included in the CRISPRi library. Over-representation of GO BP terms was assessed using Fisher’s exact test implemented in R. Enrichment results were visualized as dot plots, with dot size representing gene counts per GO term, color indicating –log10(p-value), and the x-axis denoting fold enrichment. DrugZ normalized Z-scores (NormZ values) at the mid-point and endpoint was compared for each PARP inhibitor treatment independently using scatter plots.

### Immunofluorescence microscopy

Subnuclear localization of PCNA, PARP1, LIG3, or γH2AX was examined by indirect immunofluorescence as we previously did (Yan et al., 2025). Briefly, cells grown on glass coverslips to approximately 50% confluence were washed with PBS and fixed in cold methanol (−20 °C) for 30 min. Fixed cells were incubated with primary antibodies against PARP1 (rabbit polyclonal, Cell Signaling Technology, 1:100), PCNA (mouse monoclonal, Cell Signaling Technology, 1:100), LIG3 (rabbit monoclonal), or γH2AX (rabbit monoclonal, Millipore, 1:800), RAD51 (rabbit polyclonal, Abcam, 1:500). After washing, the slide was incubated with fluorescently labeled secondary antibodies.

Images were acquired using the Observer II fluorescence microscope or Zeiss LSM 800 confocal microscope. Images were processed and quantified using ImageJ/Fiji.

### Proximity Ligation Assay (PLA)

Proximity ligation assays (PLA) were performed to detect protein–protein interactions *in situ* using the Duolink® In Situ PLA system (Sigma-Aldrich) according to the manufacturer’s instructions with minor modifications. Cells were seeded on glass coverslips and cultured to 2 × 10 cells per well in 24-well plates in the presence or absence of the PARP inhibitor olaparib, as indicated. Cells were fixed with 4% paraformaldehyde in PBS or 100% pre-cold methanol for 30 min at room temperature, followed by permeabilization with 0.2% Triton X-100 in PBS for 15 min or acetone for 1 min. After washing with PBS, nonspecific binding was blocked using Duolink blocking solution for 1 h at 37 °C. Cells were then incubated with primary antibodies raised in different species (as specified in each experiment) diluted in antibody diluent overnight at 4 °C. After washing, species-specific PLA probes (PLUS and MINUS) were applied and incubated for 1 h at 37 °C. Ligation was performed using the ligation solution for 30 min at 37 °C, followed by rolling-circle amplification with polymerase for 100 min at 37 °C. After amplification, cells were washed and counterstained with DAPI to visualize nuclei. Coverslips were mounted using Duolink mounting medium and sealed. PLA signals were visualized and recorded using the LSM 800 or LSM 900 confocal microscope. Quantification of PLA signals was performed using ImageJ/Fiji. Negative controls used omission of one primary antibody.

### Click-iT EdU Cell Proliferation Assays

To label newly synthesized DNA and assess subnuclear replication-associated signals, cells were pulse-labeled with 10 μM EdU for 20-30 min prior to fixation. Cells were subsequently fixed and permeabilized as described in the *Proximity Ligation Assay* section. Detection of incorporated EdU was carried out using a copper-catalyzed click chemistry reaction containing 2 mM CuSO₄, 10 μM Alexa Fluor 488 azide (or azide– biotin for PLA experiments), and 50 mM sodium ascorbate in PBS. The reaction was performed for 1 h at room temperature, protected from light where appropriate.

Following the click reaction, cells were washed with PBS and incubated with Image-iT™ FX Signal Enhancer (Invitrogen) for 30 min at room temperature to reduce background. Immunostaining with primary and secondary antibodies was performed as described in the *Immunofluorescence* section or *Proximity Ligation Assay (PLA)* section.

Fluorescence images were acquired using either a Zeiss Observer II fluorescence microscope or LSM 800 or LSM 900 confocal microscope under identical imaging conditions.

### In Vitro Gap-Filling and Strand-Displacement Assay

In vitro DNA synthesis assays were performed using a defined gapped DNA substrate containing a 25 nt 5′ FAM–labeled primer annealed to a 180 nt complementary template and a 150 nt downstream oligonucleotides. Reactions (10 μL) were carried out in buffer containing 20mM Tris-HCl, pH 7.5, 8mM MgOAc2, 1mM DTT, 0.1mg/mL BSA, supplemented with 400 μM each dNTP. The gapped DNA substrate was used at a final concentration of 50 nM. Where indicated, recombinant human PARP1 was preincubated with the DNA substrate in the presence or absence of NAD⁺ (1 mM) at 37 °C for 10 min to allow PARylation. DNA synthesis was initiated by addition of DNA polymerase δ (50 ng) and PCNA (50 ng), and reactions were incubated at 37 °C for 30 min. Reactions were terminated by addition of an equal volume of denaturing stop buffer (95% formamide, 20 mM EDTA) and heat-denatured at 95 °C for 5 min. Reaction products were resolved on 15% denaturing urea–polyacrylamide gels and visualized by fluorescence imaging of the FAM label. Gap-filling and strand-displacement products were identified and quantified based on electrophoretic mobility.

For the in vitro DNA synthesis assay using ³²P-dATP, reactions (10 μL) were carried out in buffer containing 20mM Tris-HCl, pH 7.5, 8mM MgOAc2, 1mM DTT, 0.1mg/mL BSA. The DNA substrate was used at a final concentration of 50 nM. Reaction mixtures were supplemented with 400 μM each dCTP, dGTP, and dTTP, and ³²P-dATP. The reactions were carried out with or without Olaparib or Veliparib. DNA synthesis was initiated by addition of DNA polymerase δ and PCNA and incubated at 37 °C for 30 min. Reactions were terminated by addition of denaturing stop buffer (95% formamide, 20 mM EDTA) and heated at 95 °C for 5 min. Products were resolved on 10% denaturing urea– polyacrylamide gels. Radiolabeled DNA products were visualized by phosphor imaging, and gap-filling and strand-displacement products were identified based on electrophoretic mobility.

### Transmission Electron Microscopy of DNA Replication Forks

DNA replication intermediates were analyzed by transmission electron microscopy (TEM) following established procedures (Zellweger and Lopes, 2018) (Shi et al., 2024) with minor modifications. To stabilize replication fork structures in vivo, cells were incubated with 10 μg/mL 4,5′,8-trimethylpsoralen (psoralen) for 5 min at 4 °C and irradiated with long-wavelength UV light (366 nm) to induce interstrand DNA crosslinking. This treatment was repeated four times. Genomic DNA was isolated under gentle conditions to preserve replication intermediates. Briefly, cell pellets were resuspended in lysis buffer containing 1.28 M Sucrose; 40 mM Tris–HCl pH 7.5; 20 mM MgCl2; 4% Triton X-100, supplemented with proteinase K, and incubated at 50 °C for 2.5 hours. RNA was removed by RNase A treatment. DNA was extracted by phenol– chloroform extraction and precipitated with 70% ethanol. Purified DNA was resuspended in TE buffer. DNA replication intermediates were enriched by benzoylated– naphthoylated DEAE-cellulose chromatography. PvuII-digested genomic DNA was loaded onto the column under low-salt conditions, and linear duplex DNA was selectively eliminated through stepwise salt elution. Single-stranded DNA–containing fractions, including replication forks and replication bubbles, were selectively eluted with a high-salt buffer containing caffeine. Eluted DNA was precipitated, resuspended, and subjected to buffer exchange to remove remaining salts and particulates before grid preparation. For electron microscopy, DNA molecules were spread onto carbon-coated copper grids using the spreading method in the presence of benzyl dimethyl alkyl ammonium chloride (BAC) as a cationic detergent to facilitate uniform DNA adsorption. Grids were air-dried and rotary shadowed with platinum–palladium at a low angle.

Samples were examined using a transmission electron microscope operated at 120 kV. Images were acquired digitally at 10,000x-20,000× magnification. Quantification was performed on randomly selected DNA molecules from at least three independent experiments.

### In situ 5’ flap labeling using D181A-FEN1

Biotin-linked D181A-FEN1 was prepared using the amine reaction-based protein biotin labeling kit and purified recombinant D181A FEN1 protein (Thermo Fisher) following the supplier’s instruction. To label 5’ flap in situ using Alexa 568-D181A, cells were seeded onto glass coverslips placed in 12-well plates, which were pre-coated with poly-L-lysine when indicated. Approximately 2-3 × 10 cells per well were seeded. To define replicating cells, the seeded cells were pulse-labeled with 10 μM EdU by replacing the medium with fresh EdU-containing medium for 20-30 min at 37 °C. EdU labeling was terminated by removing the medium and washing cells twice with warm PBS for 5 min each. Cells were fixed using either 4% paraformaldehyde (PFA) in PBS for 30 min at room temperature, followed by permeabilization with 0.2% Triton X-100 for 15 min, or with ice-cold 100% methanol at −20 °C for 30 min, in which case no additional permeabilization step was performed. Cells were washed with PBS as indicated between steps. EdU incorporation was detected using a Click-iT reaction. The reaction mixture was prepared freshly in PBS containing 2 mM CuSO₄, 10 μM Alexa Fluor 488–azide, and 50 mM sodium ascorbate. Coverslips were incubated for 1 h at room temperature protected from light, followed by three washes with PBS. Cells were blocked with Image-iT FX Signal Enhancer for 30 min at room temperature in the dark and rinsed with PBS. For D181A-FEN1 binding, a binding buffer containing 20 mM Tris-HCl (pH 8.0), 125 mM NaCl, 5 mM MgCl₂, 0.1 mg/mL BSA, and 1 mM DTT was freshly prepared and kept ice-cold. Biotylated D181A-FEN1 was diluted to 20 ng/μL in binding buffer and incubated with cells at 4 °C for 2 h. Coverslips were washed three times with cold binding buffer for 10 min each and rinsed once with PBS. Proteins were then fixed with 4% PFA for 10 min at room temperature, followed by two washes with PBS. When methanol fixation was used initially, cells were permeabilized with 0.2% Triton X-100 for 15 min at this stage. After washing with PBS, cells were incubated with Alexa Fluor 568–conjugated avidin diluted 1:100 in PBS for 1 h at room temperature. Coverslips were washed three times with PBS for 10 min each, counterstained with DAPI (1:25,000) for 5 min, and washed twice with PBS prior to mounting.

### Neutral Comet Assay

DNA double-strand breaks were assessed using a neutral comet assay performed with the CometAssay® Kit (Trevigen, 4250-050). Approximately 1 × 10 cells were mixed with molten low-melting-point agarose and immediately spread onto comet slides. Cells were lysed in neutral lysis buffer at 4 °C for 1 h to remove membranes and soluble proteins while preserving double-strand DNA integrity. Slides were then equilibrated in neutral electrophoresis buffer and subjected to electrophoresis under TBE buffer at 21 V for 45 min at 4 °C. Following electrophoresis, slides were washed with distilled water and fixed with 70% ethanol for 10 min. DNA was stained with SYBR™ Gold for 30 min and visualized using a fluorescence microscope. Comet images were analyzed using ImageJ/OpenComet software, and DNA damage was quantified as tail moment. At least 50–100 cells were analyzed per condition in each independent experiment.

### BrdU Comet Assay

DNA strand breaks in newly synthesized DNA were assessed using a BrdU-based alkaline comet assay. Cells were pulse-labeled with 20 μM BrdU for 20 min at 37 °C to mark nascent DNA. Cells were harvested, resuspended in ice-cold PBS with a concentration of approximately 1X10^5^ cells/ mL, and embedded in low–melting point agarose on comet slides. A volume of 10 µL of the cell suspension was mixed with 100 µL of 0.5% low-melting-point agarose (LMPA) at 37 °C. After agarose solidification, slides were incubated in lysis buffer (Trevigen, 4250-050) at 4 °C for 1 h, followed by alkaline unwinding in Alkaline Unwinding Solution (200mM NaOH, 1mM EDTA) at 4 °C for 1 h to unwind the DNA. Electrophoresis was carried out at 21 V for 30 minutes at 4 °C in Alkaline Electrophoresis Solution to allow separation of fragmented DNA. DNA was then washed with distilled water, fixed in 70% ethanol for 10 minutes, and allowed air dry in 37 °C. Incorporated BrdU was detected using an anti-BrdU (BD 347580) antibody, followed by a fluorescently labeled secondary antibody. Comet images were acquired using on a observe II microscope and ZEN 3.1 software. DNA damage in nascent strands was quantified by measuring tail moment using ImageJ/OpenComet software.

### Whole-Cell Lysate Preparation and Immunoblotting

Whole-cell lysates from wild-type and LIG1KD cells were prepared by washing cells with ice-cold PBS and lysing them in RIPA buffer supplemented with cocktail inhibitors.

Lysates were incubated on ice with occasional mixing and clarified by centrifugation at 14,000 × g for 15 min at 4 °C. Protein concentrations were determined using a BCA assay. Equal amounts of protein were resolved by SDS–PAGE and transferred onto PVDF membranes. Membranes were blocked with 5% nonfat dry milk or BSA in TBST and incubated with primary antibodies diluted in blocking buffer at 4 °C overnight. After washing with TBST, membranes were incubated with HRP-conjugated secondary antibodies at room temperature. Immunoreactive bands were detected using enhanced chemiluminescence and imaged using a digital imaging system.

### Cell proliferation and PARP inhibitor sensitivity analysis

Cell proliferation rate was analyzed by directly cell counting. 5 x 10^3^ cells (WT or LIG1 knockdown MDA-MB-231) were seeded in 12-well plates. Cells were grown in DMEM containing 10% FBS, 1% Penicillin-Streptomycin, and varying concentrations of PARP inhibitors [Olaparib, Talazoparib (BMN-673), and Veliparib (ABT-888)]. The culture medium was replaced with fresh medium every other day. On day 7, the live cell number was counted using a hemocytometer. 5 x 10^3^ cells were re-seeded in 12 well plates and grown in the same medium as in day 0-day 7. On day 14, the live cell number was counted using a hemocytometer.

To measure the PARP inhibition sensitivity, 1 × 10 WT or LIG1 knockdown MDA-MB- 231 cells into 12-well plates. Cells were grown in DMEM containing 10% FBS, 1% Penicillin-Streptomycin, and varying concentrations of PARP inhibitors for 4 days. Cell viability was assessed by counting viable cells, and survival rates were expressed relative to untreated controls.

### Whole-exome sequencing (WES) and data analysis

WES were conducted on genomic DNA isolated from WT or LIG1 knockdown cells with or without olaparib treatment (4 days). WES library preparation and DNA sequencing were carried out as we previously described (Yan et al., 2025). WES sequencing reads were assessed using FastQC and filtered to remove the adaptor sequence. We selected the paired end reads that were longer than 35 bp after trimming for analysis. High-quality paired end reads from each sample were separately aligned to the reference mouse genome mm10 using Bowtie2 (Langmead and Salzberg, 2012). The binary alignment map was sorted and indexed using Samtools and Read duplicates were removed from the sorted BAM file to create a BAM file with unique reads only.

Single nucleotide variations or small insertions or deletions were analyzed using the VarScan software package (Koboldt et al., 2012). Germline mutations were filtered out by setting the allele frequency in the WT control as 0. The somatic mutations were scored with the mutant allele frequency in the sample no less than 0.05. Tandem duplications were analyzed using Pindel (Ye et al., 2009). Somatic duplications were scored if at least three supporting tracks in the sample were detected but no supporting tracks were detected in the WT control. The frequencies of somatic mutations or duplications were calculated by dividing the number of somatic events by estimated size of mouse exome (∼30 millions). We used the SigProfilerClusters program (Bergstrom et al., 2022) to define mutation clusters. A window size of 1 Mb was used to adjust intra-mutational distances based on local mutation density and variant allele frequencies as well. The mutation signature analysis was carried out using the SigProfilerAssignment program (Díaz-Gay et al., 2023).

**Extended Data can be found on the *Nature Structural & Molecular Biology* website.**

## Acknowledgements

We thank Light Microscopy Digital Imaging (LMDI) Shared Resource at City of Hope (COH) for assistance with IF imaging analysis. Research reported in this publication included work performed in the LMDI and the Integrate Genomics and Bioinformatics Shared Resources at COH was supported by the National Cancer Institute of the National Institutes of Health under grant number P30 CA033572. The content is solely the responsibility of the authors and does not necessarily represent the official views of the National Institutes of Health. This work was supported by NIH grants R50 CA211397 to L.Z. and R01 CA073764 to B.S.

## Author information

The authors declare there are no conflicts of interest in this study. Correspondence and requests for materials should be addressed to B.S..

