## Supplemental for "PARP1-mediated 5’ flap dynamics facilitate Okazaki fragment maturation"

### Slide 1
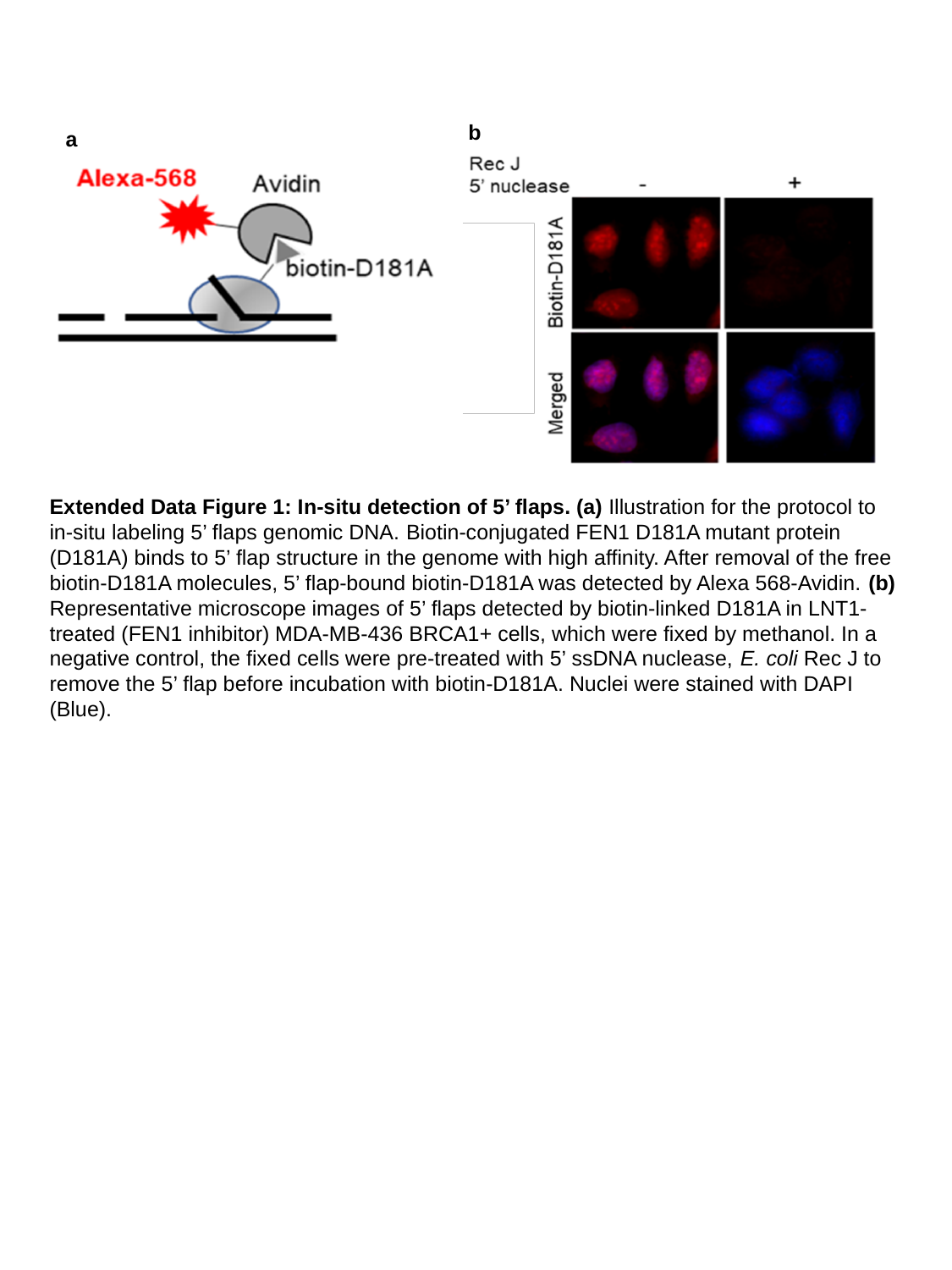

b
a
Extended Data Figure 1: In-situ detection of 5’ flaps. (a) Illustration for the protocol to in-situ labeling 5’ flaps genomic DNA. Biotin-conjugated FEN1 D181A mutant protein (D181A) binds to 5’ flap structure in the genome with high affinity. After removal of the free biotin-D181A molecules, 5’ flap-bound biotin-D181A was detected by Alexa 568-Avidin. (b) Representative microscope images of 5’ flaps detected by biotin-linked D181A in LNT1-treated (FEN1 inhibitor) MDA-MB-436 BRCA1+ cells, which were fixed by methanol. In a negative control, the fixed cells were pre-treated with 5’ ssDNA nuclease, E. coli Rec J to remove the 5’ flap before incubation with biotin-D181A. Nuclei were stained with DAPI (Blue).

### Slide 2
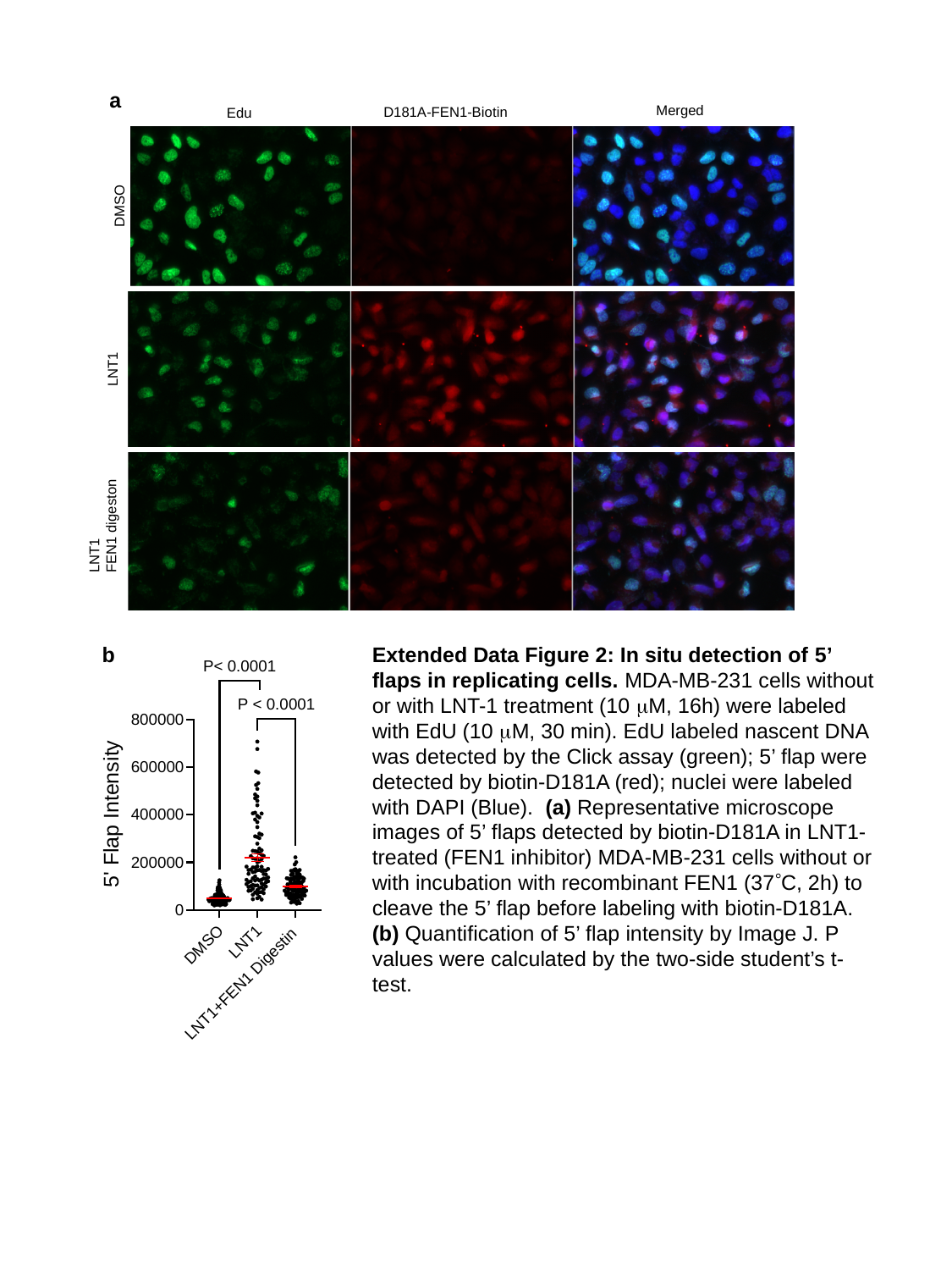

a
Merged
D181A-FEN1-Biotin
Edu
DMSO
LNT1
LNT1
FEN1 digeston
b
Extended Data Figure 2: In situ detection of 5’ flaps in replicating cells. MDA-MB-231 cells without or with LNT-1 treatment (10 μM, 16h) were labeled with EdU (10 μM, 30 min). EdU labeled nascent DNA was detected by the Click assay (green); 5’ flap were detected by biotin-D181A (red); nuclei were labeled with DAPI (Blue). (a) Representative microscope images of 5’ flaps detected by biotin-D181A in LNT1-treated (FEN1 inhibitor) MDA-MB-231 cells without or with incubation with recombinant FEN1 (37°C, 2h) to cleave the 5’ flap before labeling with biotin-D181A. (b) Quantification of 5’ flap intensity by Image J. P values were calculated by the two-side student’s t-test.

### Slide 3
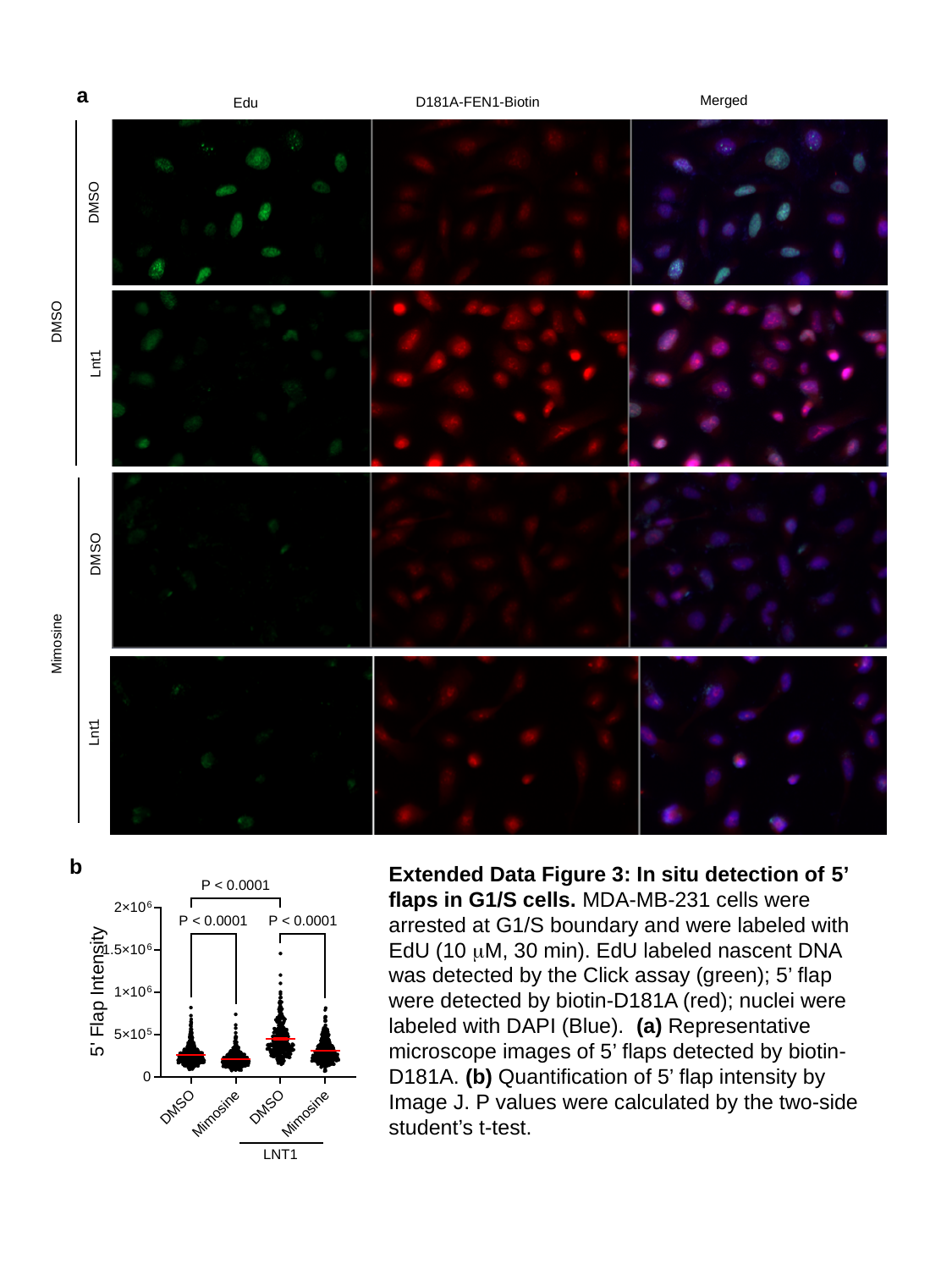

a
Merged
D181A-FEN1-Biotin
Edu
DMSO
DMSO
Lnt1
DMSO
Mimosine
Lnt1
b
Extended Data Figure 3: In situ detection of 5’ flaps in G1/S cells. MDA-MB-231 cells were arrested at G1/S boundary and were labeled with EdU (10 μM, 30 min). EdU labeled nascent DNA was detected by the Click assay (green); 5’ flap were detected by biotin-D181A (red); nuclei were labeled with DAPI (Blue). (a) Representative microscope images of 5’ flaps detected by biotin-D181A. (b) Quantification of 5’ flap intensity by Image J. P values were calculated by the two-side student’s t-test.

### Slide 4
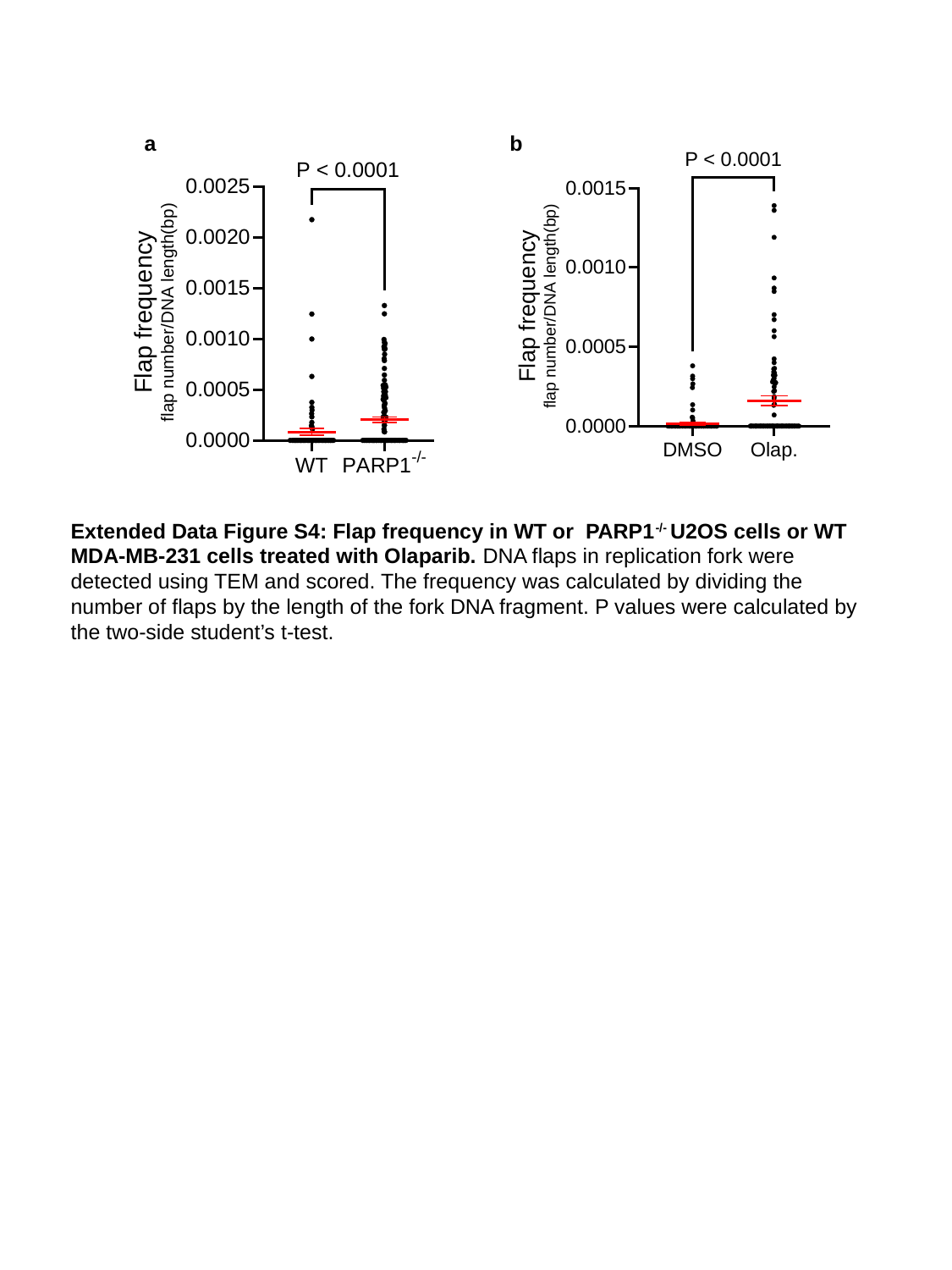

a
b
Extended Data Figure S4: Flap frequency in WT or PARP1^-/-^ U2OS cells or WT MDA-MB-231 cells treated with Olaparib. DNA flaps in replication fork were detected using TEM and scored. The frequency was calculated by dividing the number of flaps by the length of the fork DNA fragment. P values were calculated by the two-side student’s t-test.

### Slide 5
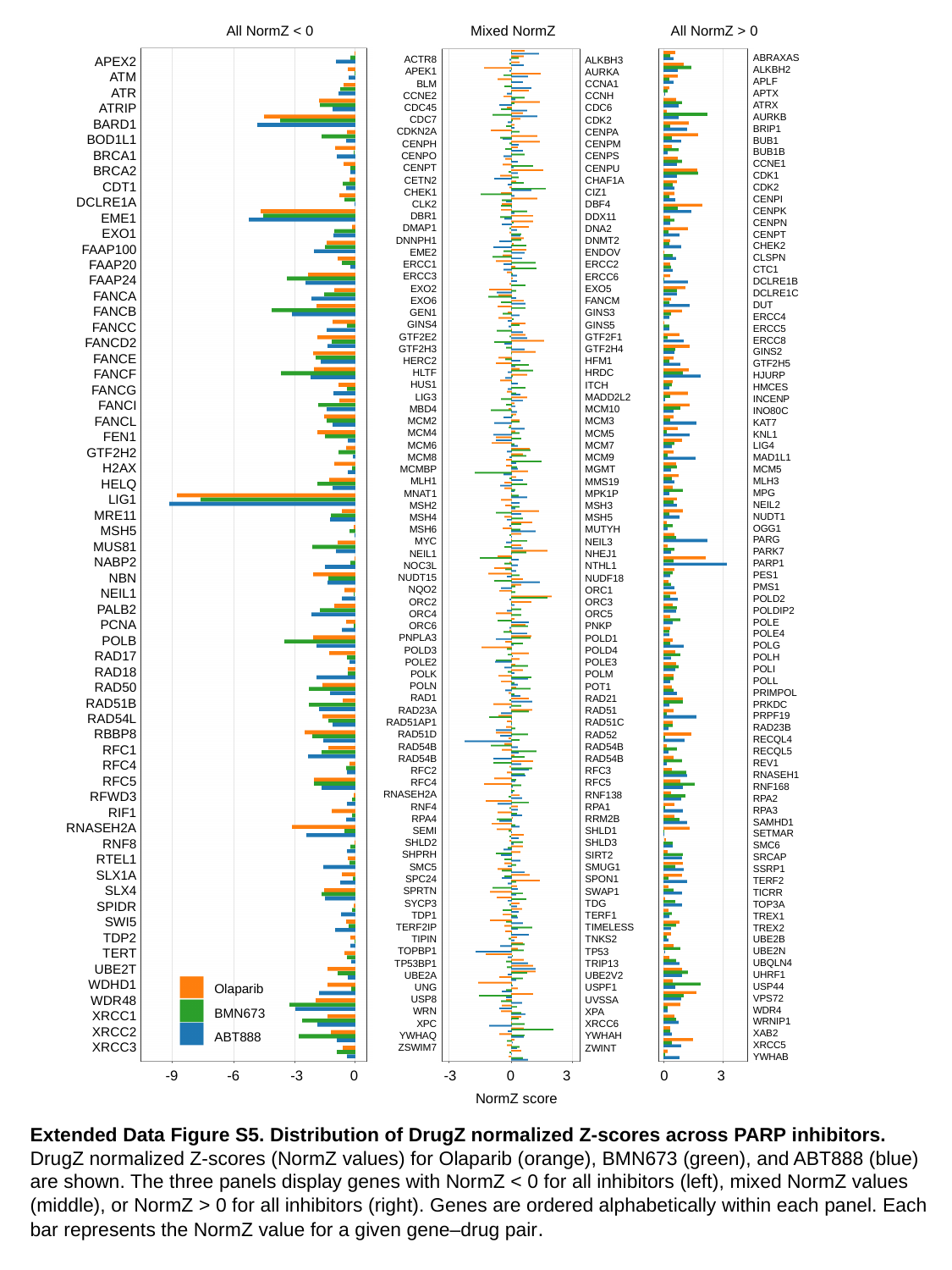

All NormZ < 0
Mixed NormZ
All NormZ > 0
ABRAXAS
ALKBH2
APLF
APTX
ATRX
AURKB
BRIP1
BUB1
BUB1B
CCNE1
CDK1
CDK2
CENPI
CENPK
CENPN
CENPT
CHEK2
CLSPN
CTC1
DCLRE1B
DCLRE1C
DUT
ERCC4
ERCC5
ERCC8
GINS2
GTF2H5
HJURP
HMCES
INCENP
INO80C
KAT7
KNL1
LIG4
MAD1L1
MCM5
MLH3
MPG
NEIL2
NUDT1
OGG1
PARG
PARK7
PARP1
PES1
PMS1
POLD2
POLDIP2
POLE
POLE4
POLG
POLH
POLI
POLL
PRIMPOL
PRKDC
PRPF19
RAD23B
RECQL4
RECQL5
REV1
RNASEH1
RNF168
RPA2
RPA3
SAMHD1
SETMAR
SMC6
SRCAP
SSRP1
TERF2
TICRR
TOP3A
TREX1
TREX2
UBE2B
UBE2N
UBQLN4
UHRF1
USP44
VPS72
WDR4
WRNIP1
XAB2
XRCC5
YWHAB
APEX2
ATM
ATR
ATRIP
BARD1
BOD1L1
BRCA1
BRCA2
CDT1
DCLRE1A
EME1
EXO1
FAAP100
FAAP20
FAAP24
FANCA
FANCB
FANCC
FANCD2
FANCE
FANCF
FANCG
FANCI
FANCL
FEN1
GTF2H2
H2AX
HELQ
LIG1
MRE11
MSH5
MUS81
NABP2
NBN
NEIL1
PALB2
PCNA
POLB
RAD17
RAD18
RAD50
RAD51B
RAD54L
RBBP8
RFC1
RFC4
RFC5
RFWD3
RIF1
RNASEH2A
RNF8
RTEL1
SLX1A
SLX4
SPIDR
SWI5
TDP2
TERT
UBE2T
WDHD1
WDR48
XRCC1
XRCC2
XRCC3
ACTR8
APEK1
BLM
CCNE2
CDC45
CDC7
CDKN2A
CENPH
CENPO
CENPT
CETN2
CHEK1
CLK2
DBR1
DMAP1
DNNPH1
EME2
ERCC1
ERCC3
EXO2
EXO6
GEN1
GINS4
GTF2E2
GTF2H3
HERC2
HLTF
HUS1
LIG3
MBD4
MCM2
MCM4
MCM6
MCM8
MCMBP
MLH1
MNAT1
MSH2
MSH4
MSH6
MYC
NEIL1
NOC3L
NUDT15
NQO2
ORC2
ORC4
ORC6
PNPLA3
POLD3
POLE2
POLK
POLN
RAD1
RAD23A
RAD51AP1
RAD51D
RAD54B
RAD54B
RFC2
RFC4
RNASEH2A
RNF4
RPA4
SEMI
SHLD2
SHPRH
SMC5
SPC24
SPRTN
SYCP3
TDP1
TERF2IP
TIPIN
TOPBP1
TP53BP1
UBE2A
UNG
USP8
WRN
XPC
YWHAQ
ZSWIM7
ALKBH3
AURKA
CCNA1
CCNH
CDC6
CDK2
CENPA
CENPM
CENPS
CENPU
CHAF1A
CIZ1
DBF4
DDX11
DNA2
DNMT2
ENDOV
ERCC2
ERCC6
EXO5
FANCM
GINS3
GINS5
GTF2F1
GTF2H4
HFM1
HRDC
ITCH
MADD2L2
MCM10
MCM3
MCM5
MCM7
MCM9
MGMT
MMS19
MPK1P
MSH3
MSH5
MUTYH
NEIL3
NHEJ1
NTHL1
NUDF18
ORC1
ORC3
ORC5
PNKP
POLD1
POLD4
POLE3
POLM
POT1
RAD21
RAD51
RAD51C
RAD52
RAD54B
RAD54B
RFC3
RFC5
RNF138
RPA1
RRM2B
SHLD1
SHLD3
SIRT2
SMUG1
SPON1
SWAP1
TDG
TERF1
TIMELESS
TNKS2
TP53
TRIP13
UBE2V2
USPF1
UVSSA
XPA
XRCC6
YWHAH
ZWINT
Olaparib
BMN673
ABT888
-9
-6
-3
0
-3
0
3
0
3
NormZ score
Extended Data Figure S5. Distribution of DrugZ normalized Z-scores across PARP inhibitors. DrugZ normalized Z-scores (NormZ values) for Olaparib (orange), BMN673 (green), and ABT888 (blue) are shown. The three panels display genes with NormZ < 0 for all inhibitors (left), mixed NormZ values (middle), or NormZ > 0 for all inhibitors (right). Genes are ordered alphabetically within each panel. Each bar represents the NormZ value for a given gene–drug pair.

### Slide 6
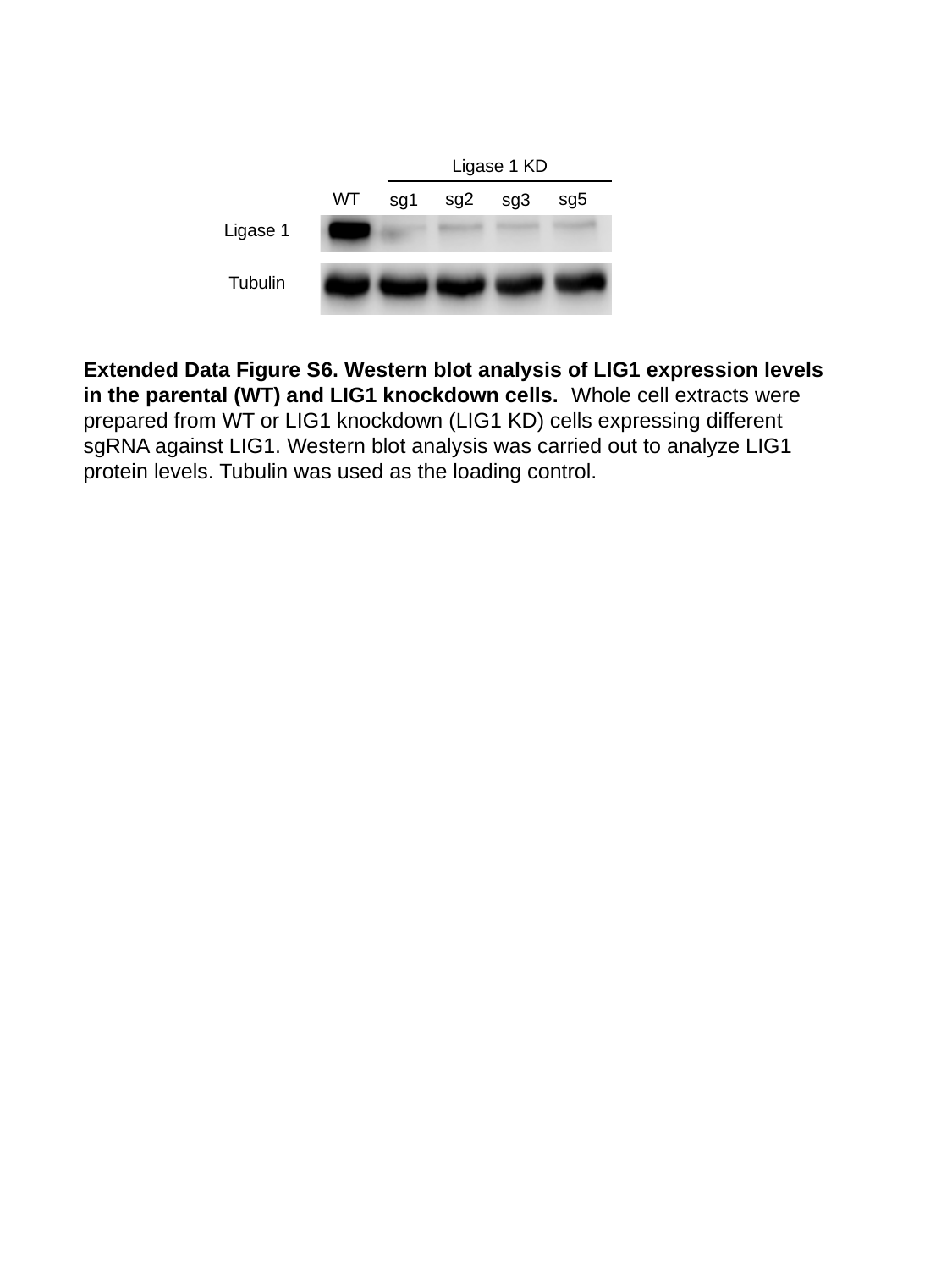

Ligase 1 KD
sg5
sg2
WT
sg1
sg3
Ligase 1
Tubulin
Extended Data Figure S6. Western blot analysis of LIG1 expression levels in the parental (WT) and LIG1 knockdown cells. Whole cell extracts were prepared from WT or LIG1 knockdown (LIG1 KD) cells expressing different sgRNA against LIG1. Western blot analysis was carried out to analyze LIG1 protein levels. Tubulin was used as the loading control.

### Slide 7
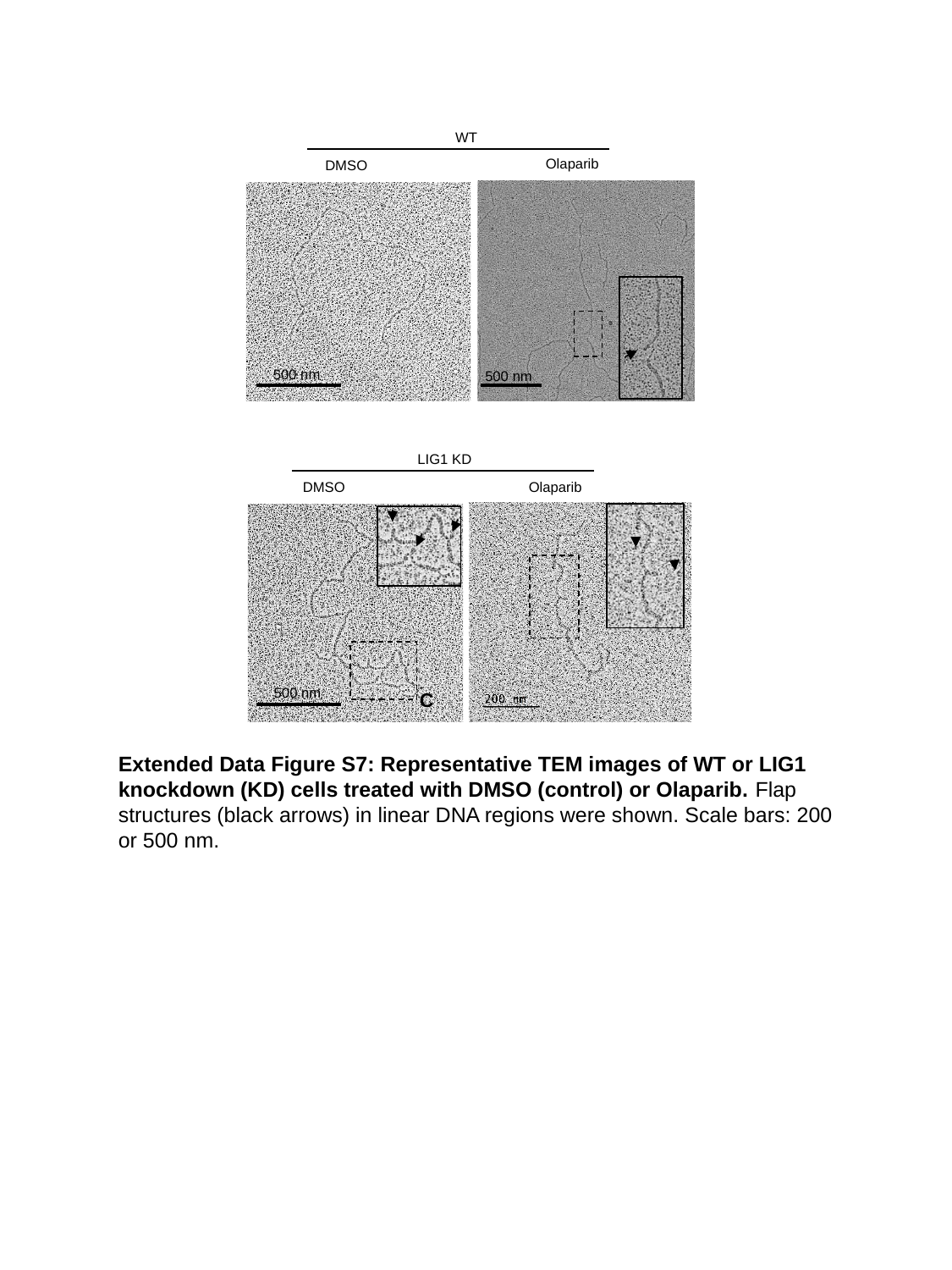

WT
Olaparib
DMSO
500 nm
500 nm
LIG1 KD
DMSO
Olaparib
C
500 nm
Extended Data Figure S7: Representative TEM images of WT or LIG1 knockdown (KD) cells treated with DMSO (control) or Olaparib. Flap structures (black arrows) in linear DNA regions were shown. Scale bars: 200 or 500 nm.

### Slide 8
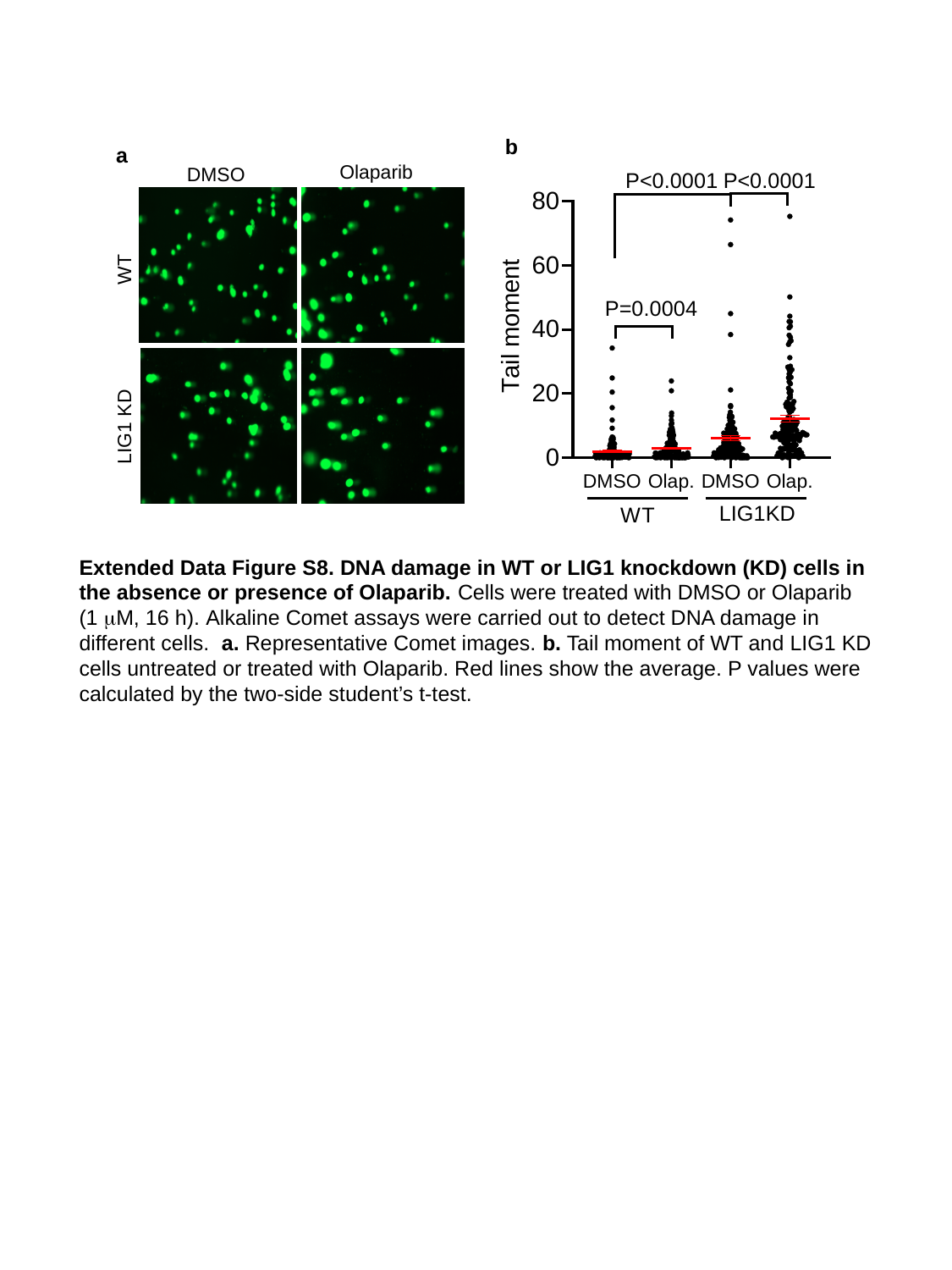

b
a
Olaparib
DMSO
WT
LIG1 KD
Extended Data Figure S8. DNA damage in WT or LIG1 knockdown (KD) cells in the absence or presence of Olaparib. Cells were treated with DMSO or Olaparib (1 μM, 16 h). Alkaline Comet assays were carried out to detect DNA damage in different cells. a. Representative Comet images. b. Tail moment of WT and LIG1 KD cells untreated or treated with Olaparib. Red lines show the average. P values were calculated by the two-side student’s t-test.

### Slide 9
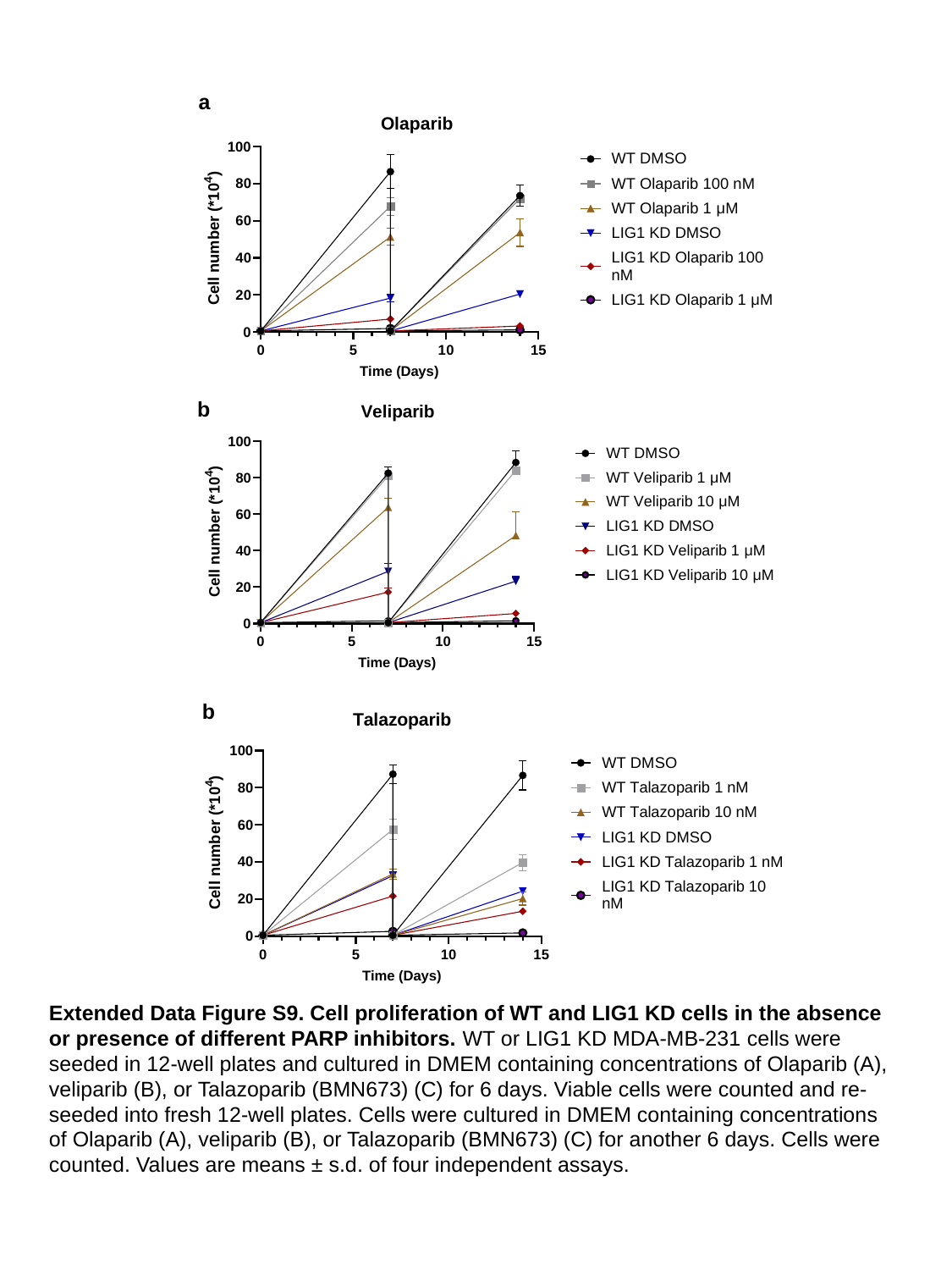

a
b
b
Extended Data Figure S9. Cell proliferation of WT and LIG1 KD cells in the absence or presence of different PARP inhibitors. WT or LIG1 KD MDA-MB-231 cells were seeded in 12-well plates and cultured in DMEM containing concentrations of Olaparib (A), veliparib (B), or Talazoparib (BMN673) (C) for 6 days. Viable cells were counted and re-seeded into fresh 12-well plates. Cells were cultured in DMEM containing concentrations of Olaparib (A), veliparib (B), or Talazoparib (BMN673) (C) for another 6 days. Cells were counted. Values are means ± s.d. of four independent assays.

### Slide 10
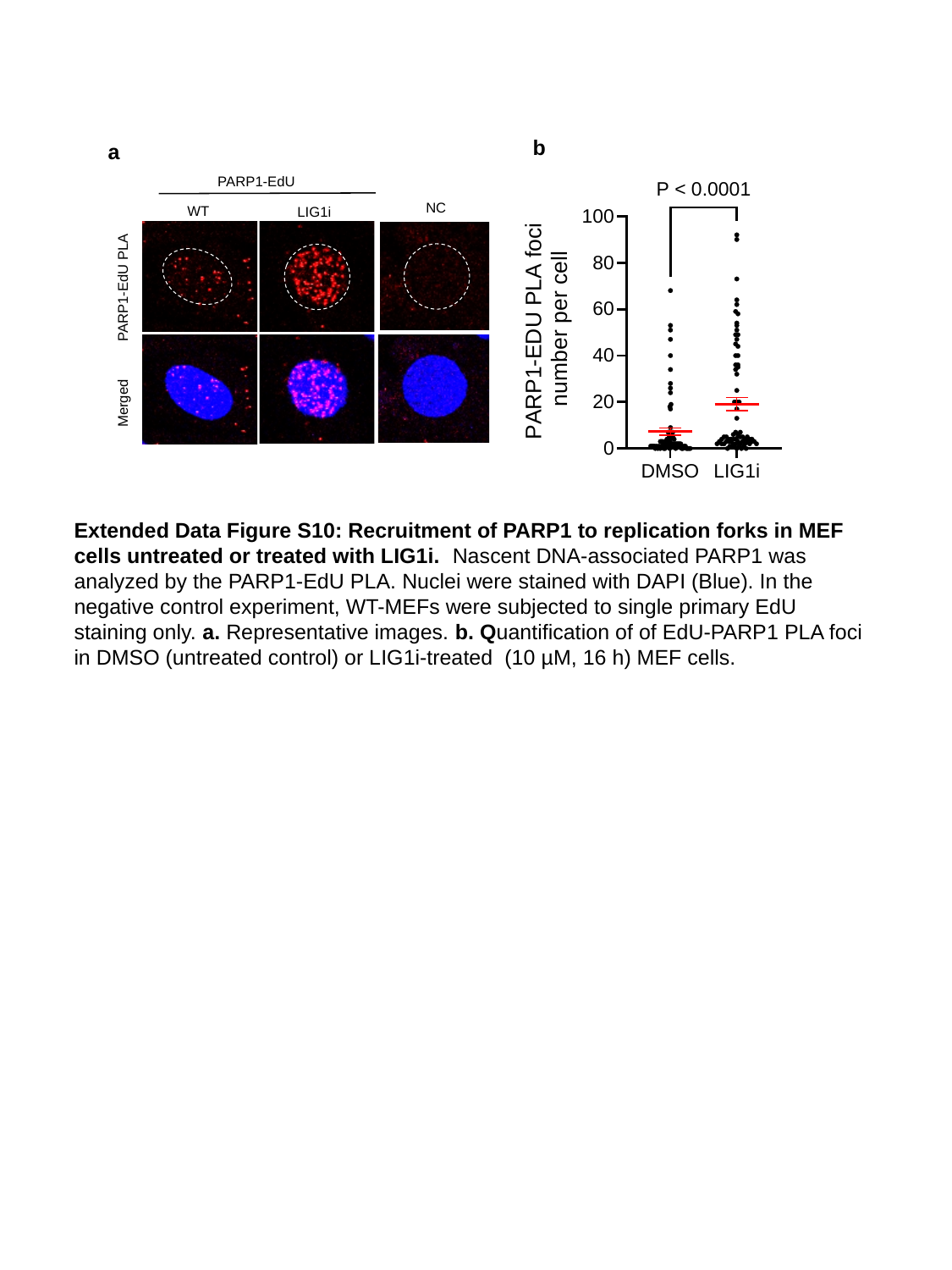

b
a
PARP1-EdU
NC
WT
LIG1i
PARP1-EdU PLA
Merged
Extended Data Figure S10: Recruitment of PARP1 to replication forks in MEF cells untreated or treated with LIG1i. Nascent DNA-associated PARP1 was analyzed by the PARP1-EdU PLA. Nuclei were stained with DAPI (Blue). In the negative control experiment, WT-MEFs were subjected to single primary EdU staining only. a. Representative images. b. Quantification of of EdU-PARP1 PLA foci in DMSO (untreated control) or LIG1i-treated (10 µM, 16 h) MEF cells.

### Slide 11
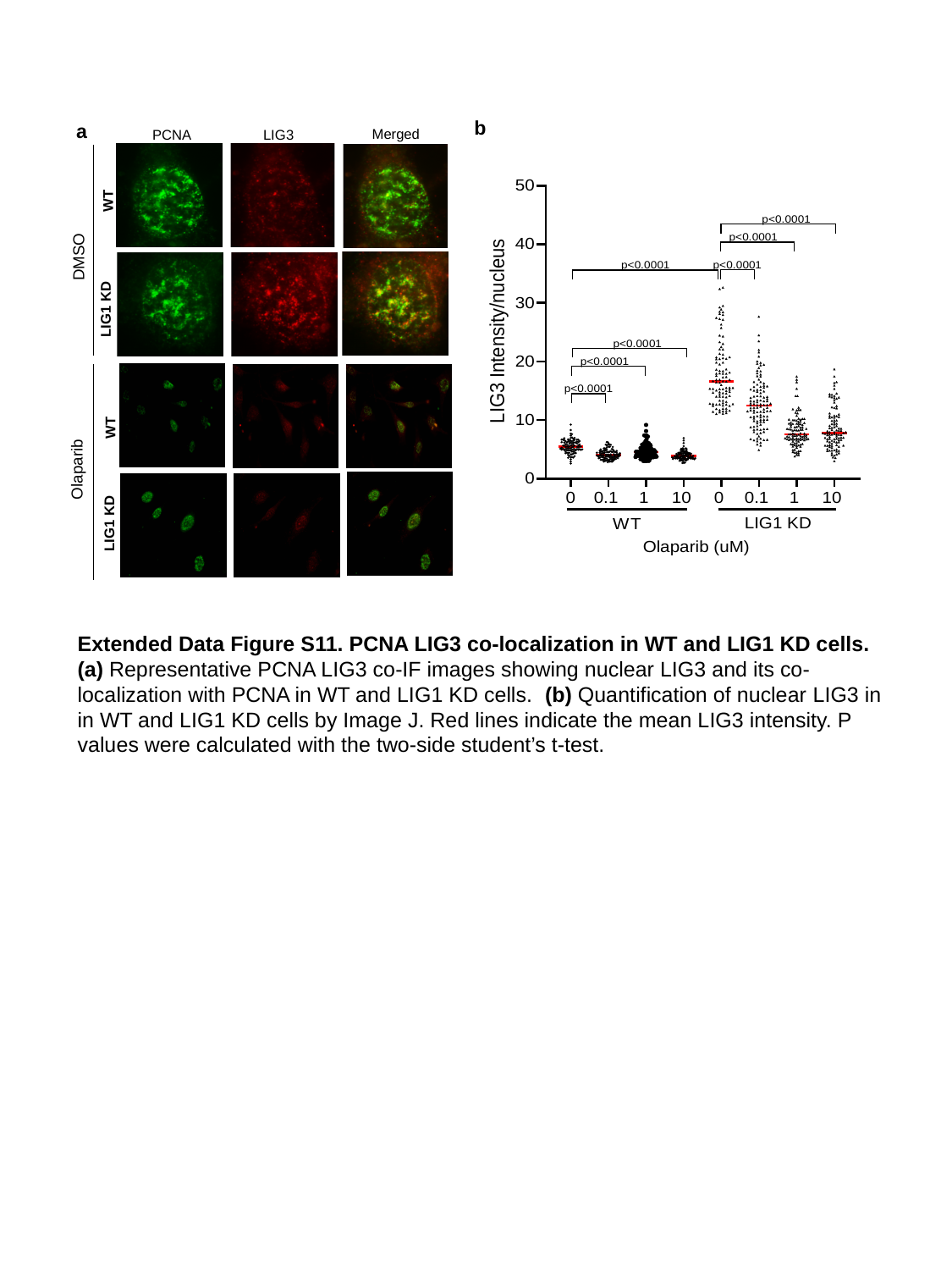

b
a
Merged
PCNA
LIG3
WT
DMSO
LIG1 KD
WT
 Olaparib
LIG1 KD
Extended Data Figure S11. PCNA LIG3 co-localization in WT and LIG1 KD cells. (a) Representative PCNA LIG3 co-IF images showing nuclear LIG3 and its co-localization with PCNA in WT and LIG1 KD cells. (b) Quantification of nuclear LIG3 in in WT and LIG1 KD cells by Image J. Red lines indicate the mean LIG3 intensity. P values were calculated with the two-side student’s t-test.

### Slide 12
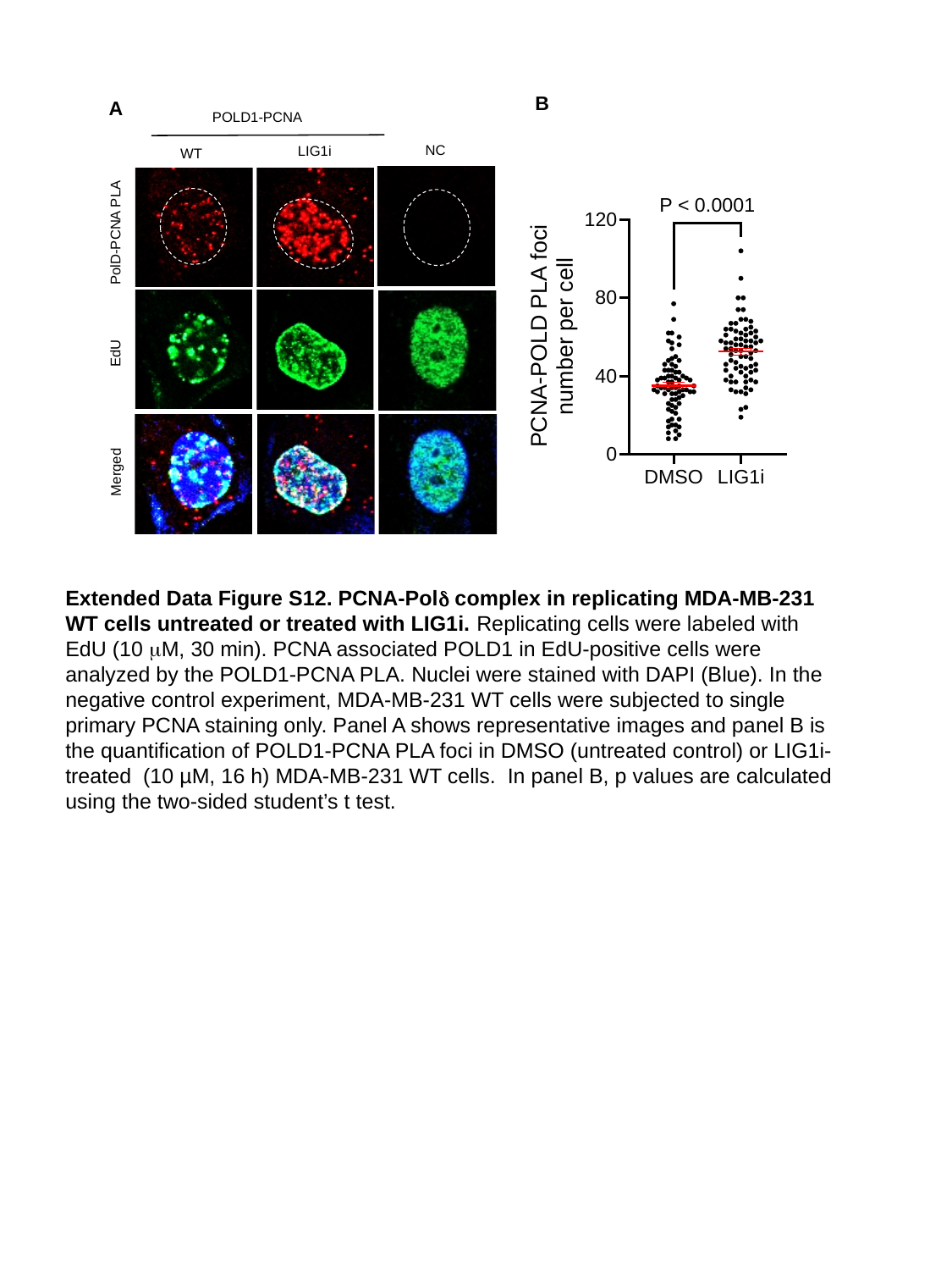

B
A
POLD1-PCNA
NC
LIG1i
WT
PolD-PCNA PLA
EdU
Merged
Extended Data Figure S12. PCNA-Polδ complex in replicating MDA-MB-231 WT cells untreated or treated with LIG1i. Replicating cells were labeled with EdU (10 μM, 30 min). PCNA associated POLD1 in EdU-positive cells were analyzed by the POLD1-PCNA PLA. Nuclei were stained with DAPI (Blue). In the negative control experiment, MDA-MB-231 WT cells were subjected to single primary PCNA staining only. Panel A shows representative images and panel B is the quantification of POLD1-PCNA PLA foci in DMSO (untreated control) or LIG1i-treated (10 µM, 16 h) MDA-MB-231 WT cells. In panel B, p values are calculated using the two-sided student’s t test.

### Slide 13
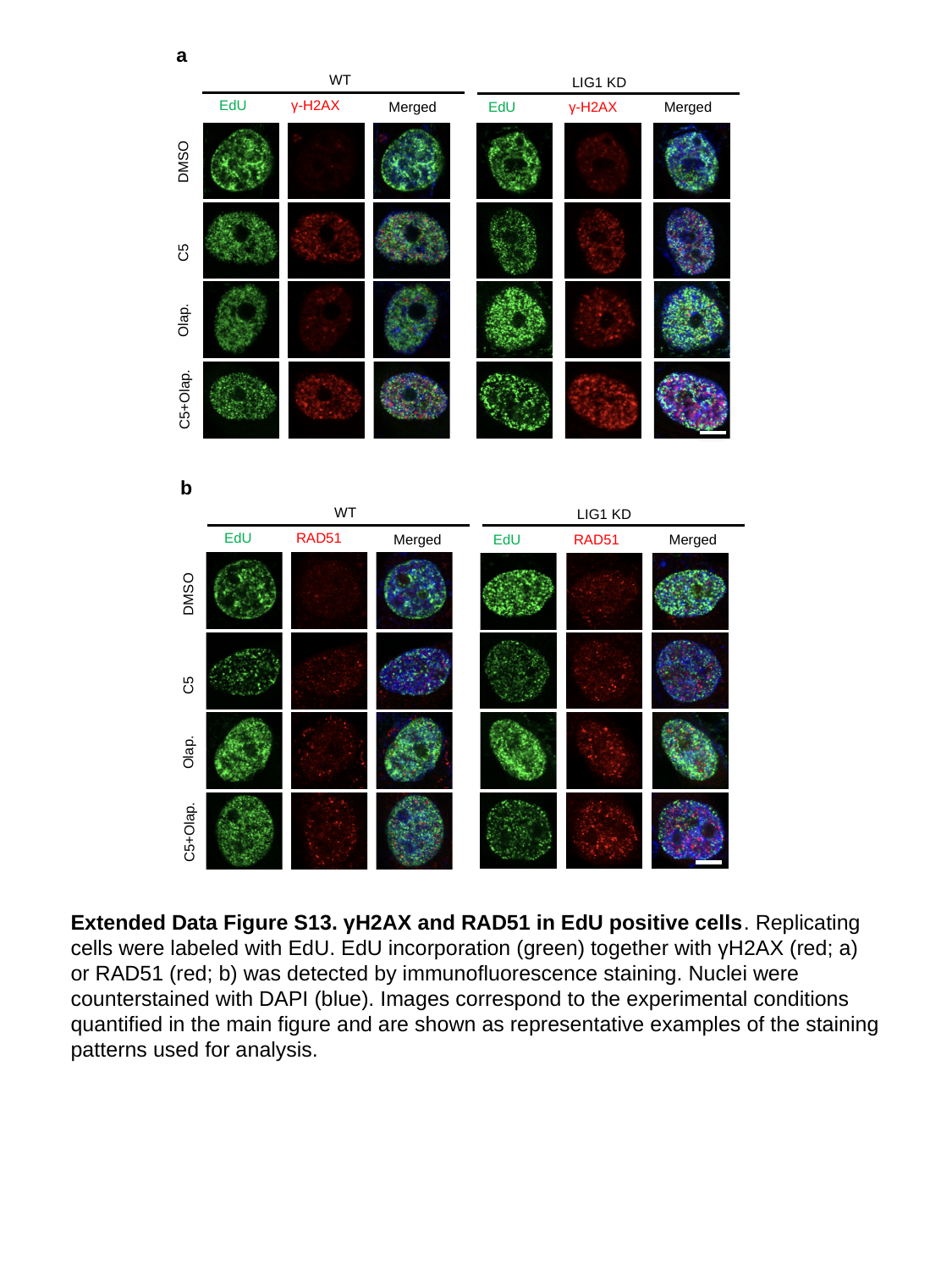

a
WT
LIG1 KD
EdU
γ-H2AX
Merged
EdU
γ-H2AX
Merged
DMSO
C5
Olap.
C5+Olap.
b
WT
LIG1 KD
EdU
RAD51
Merged
EdU
RAD51
Merged
DMSO
C5
Olap.
C5+Olap.
Extended Data Figure S13. γH2AX and RAD51 in EdU positive cells. Replicating cells were labeled with EdU. EdU incorporation (green) together with γH2AX (red; a) or RAD51 (red; b) was detected by immunofluorescence staining. Nuclei were counterstained with DAPI (blue). Images correspond to the experimental conditions quantified in the main figure and are shown as representative examples of the staining patterns used for analysis.

### Slide 14
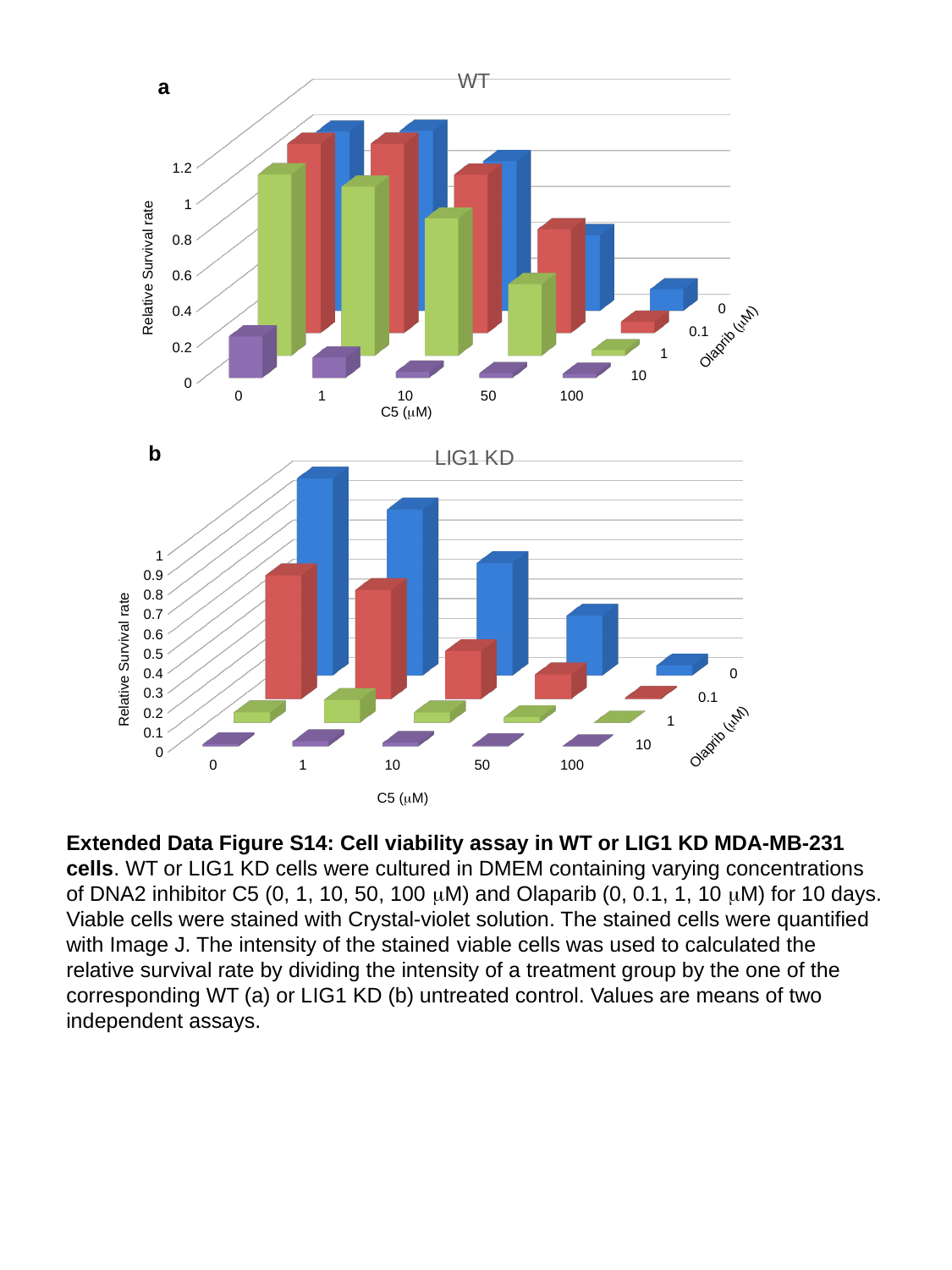

[unsupported chart]
a
Relative Survival rate
Olaprib (μM)
C5 (μM)
[unsupported chart]
b
Relative Survival rate
Olaprib (μM)
C5 (μM)
Extended Data Figure S14: Cell viability assay in WT or LIG1 KD MDA-MB-231 cells. WT or LIG1 KD cells were cultured in DMEM containing varying concentrations of DNA2 inhibitor C5 (0, 1, 10, 50, 100 μM) and Olaparib (0, 0.1, 1, 10 μM) for 10 days. Viable cells were stained with Crystal-violet solution. The stained cells were quantified with Image J. The intensity of the stained viable cells was used to calculated the relative survival rate by dividing the intensity of a treatment group by the one of the corresponding WT (a) or LIG1 KD (b) untreated control. Values are means of two independent assays.
